# Spatial proteomics of ER tubules reveals CLMN, an ER-actin tether at focal adhesions that promotes cell migration

**DOI:** 10.1101/2024.01.24.577043

**Authors:** Holly Merta, Tadamoto Isogai, Blessy Paul, Gaudenz Danuser, W Mike Henne

**Affiliations:** Department of Cell Biology, UT Southwestern Medical Center, Dallas TX 75390; Lyda Hill Department of Bioinformatics and Cecil H. and Ida Green Center for Systems Biology, UT Southwestern Medical Center, Dallas TX 75390

**Keywords:** endoplasmic reticulum (ER), TurboID, Calmin, CLMN, cellular adhesion, migration

## Abstract

The endoplasmic reticulum (ER) is structurally and functionally diverse, yet how its functions are organized within morphological subdomains is incompletely understood. Utilizing TurboID-based proximity labeling and CRISPR knock-in technologies, here we map the proteomic landscape of the human ER and nuclear envelope. Spatial proteomics reveals enrichments of proteins into ER tubules, sheets, and nuclear envelope. We uncover an ER-enriched actin-binding protein, Calmin (CLMN), and define it as an ER-actin tether that localizes to focal adhesions adjacent to ER tubules. CLMN depletion perturbs focal adhesion disassembly, actin dynamics, and cell movement. Mechanistically, CLMN-depleted cells also exhibit defects in calcium signaling near ER-actin interfaces, suggesting CLMN promotes calcium signaling near adhesions to facilitate their disassembly. Collectively, we map the sub-organelle proteome landscape of the ER, identify CLMN as an ER-actin tether, and describe a non-canonical mechanism by which ER tubules engage actin to regulate cell migration.

## Introduction

A defining feature of eukaryotes is their ability to compartmentalize biological processes into distinct organelles. Organelles themselves exhibit sub-compartmentalization, and these sub-regions adopt specific morphologies that serve as enrichment zones for proteins or complexes.

One example is the endoplasmic reticulum (ER), a vast membrane network of sheets, tubules and the nuclear envelope with diverse functions to match its structural complexity^1,2^. Whereas ER sheets act as docking sites for ribosomes and influence protein translation, ER tubules manifest other roles such as forming contacts with other organelles to exchange lipids or ions^3,4^, and to spatially coordinate mitochondrial or endosome division^5,6^. ER architecture also changes in response to metabolic cues such as nutrient stimulation or fasting, implying these morphologies support metabolic adaptation^7^. However, how the ER creates and maintains its architecture—and how its functions are spatially organized by morphological regions—remains incompletely understood. So far, evidence has been acquired to suggest that distinct morphological zones maintain their form via ER shaping proteins including the tubule-shaping reticulons (RTNs) and sheet-enriched CLIMP63^2,8,9^. ER morphology is also influenced by contacts with the cytoskeleton^1^. For example, the nuclear envelope is studded with Nesprins that tether it to actin or intermediate filaments and govern nuclear positioning^10^. In the ER periphery, ER tubules can attach to microtubules via the Tip Attachment Complex (TAC) that enables them to extend along the microtubule network^11–14^. However, whereas ER-microtubule contacts are relatively well studied, whether ER tubules interact directly also with the actin cytoskeleton remains unknown.

The ER adopts specific morphologies to support its diverse functions, and its subregions exhibit local enrichments of some proteins to facilitate this. For example, classic work on mitochondria-associated membranes (MAMs) reveal that ER-mitochondrial contacts function as subdomains that enrich enzymes important for phospholipid biosynthesis^15^. ‘Rough ER’ sheets are studded with ribosomes that facilitate translation^16^. Indeed, biochemical purification of smooth and rough ER microsomal fractions coupled with proteomics reveals distinct proteomes exist within these two ER subdomains that impact specific functions for each subregion^17^. Recent proximity-based biotinylation technologies such as APEX and TurboID^18,19^ further enable the quantitative cataloging of such proteome compartmentalization at the organelle and sub-organelle length-scales. Such proximity-based approaches have revealed high-confidence proteomes for mitochondria^20^ and for the endolysosomal pathway^21^, but a comprehensive cataloging of ER morphological subregions has not been achieved.

Here, we utilize TurboID-based proximity biotinylation combined with CRISPR-Cas9 genetic modification to catalog the local proteomic landscape of distinct ER morphological sub-domains of human cells. Equipped with this approach, we detected Calmin (CLMN) as enriched in the ER peripheral tubule network. CLMN is a poorly understood calponin homology (CH) domain-containing protein that potentially binds actin directly through its CH domains, and localizes to an unknown intracellular compartment with its transmembrane domain^22^. Beyond mapping its basic domain architecture, CLMN has not been functionally characterized. We deorphanize CLMN and mechanistically define it as a previously unappreciated ER-to-actin tether, finding it facilitates ER tubule contact with surface adhesion actin structures and influences cell migration. CLMN localizes to ER tubules via its transmembrane domain, and associates with actin via its CH domains, thus acting as the first *bona fide* ER tubule-to-actin tether. CLMN associates with actin-and paxillin-positive surface adhesions, and its loss perturbs adhesion dynamics at these sites. CLMN-deficient cells manifest altered F-actin architectures, defects in surface adhesion disassembly, and defective cell migration. We also find that CLMN-depleted cells manifest alterations in local calcium dynamics at adhesion sites. Collectively, we propose that CLMN is an ER tubule-to-actin tether that mediates ER crosstalk with surface adhesions and coordinates their disassembly during cell migration.

## Results

### TurboID-based spatial proteomics detects the protein Calmin (CLMN) in human cells as an ER tubule-enriched protein

Proteomic analyses of distinct ER membrane fractions have revealed enrichments of specific proteins partitioned by ER morphology or function. For example, rough and smooth rat liver ER-derived microsomes display only partially overlapping proteomes and feature distinct enrichments of some proteins in each fraction^17^. Similarly, reticulon-associated ER membranes derived from budding yeast display subsets of ER proteins specifically enriched in ER tubules^23^. At the same time, it is not known what functions of the ER are partitioned into ER tubules compared to sheets or the nuclear envelope (NE) in human cells. To identify proteins associated with ER morphological zones in human cells, we developed a CRISPR-Cas9 based spatial proteomics approach to profile protein distributions within the ER network of U2OS cells, which features a well-defined ER tubular and sheet network continuous with the NE^24^. We took advantage of the highly sensitive proximity-labeling biotin ligase TurboID^19^ coupled with endogenous CRISPR-Cas9 knock-in tagging^25,26^ of ER proteins known to partition into morphologically distinct ER subdomains. Five such ‘Marker Proteins’ with established ER morphological subdomain enrichments were selected and endogenously tagged with the fluorophore mNeonGreen together with TurboID: *RTN4*/reticulon 4 (ER tubules), *CKAP4*/CLIMP-63 (ER sheets), *CANX*/calnexin (general ER and NE marker), *SYNE4*/Nesprin 4 (outer NE), and *LBR*/lamin B receptor (inner NE) (Figures 1A-1B)^2,27–29^. All Marker Proteins were topologically labeled so mNeonGreen-TurboID oriented into the cytoplasm or nucleoplasm, and CRISPR knock-in cell lines were confirmed by Western blotting and confocal microscopy (Figures 1B, S1A).

**Figure 1:**
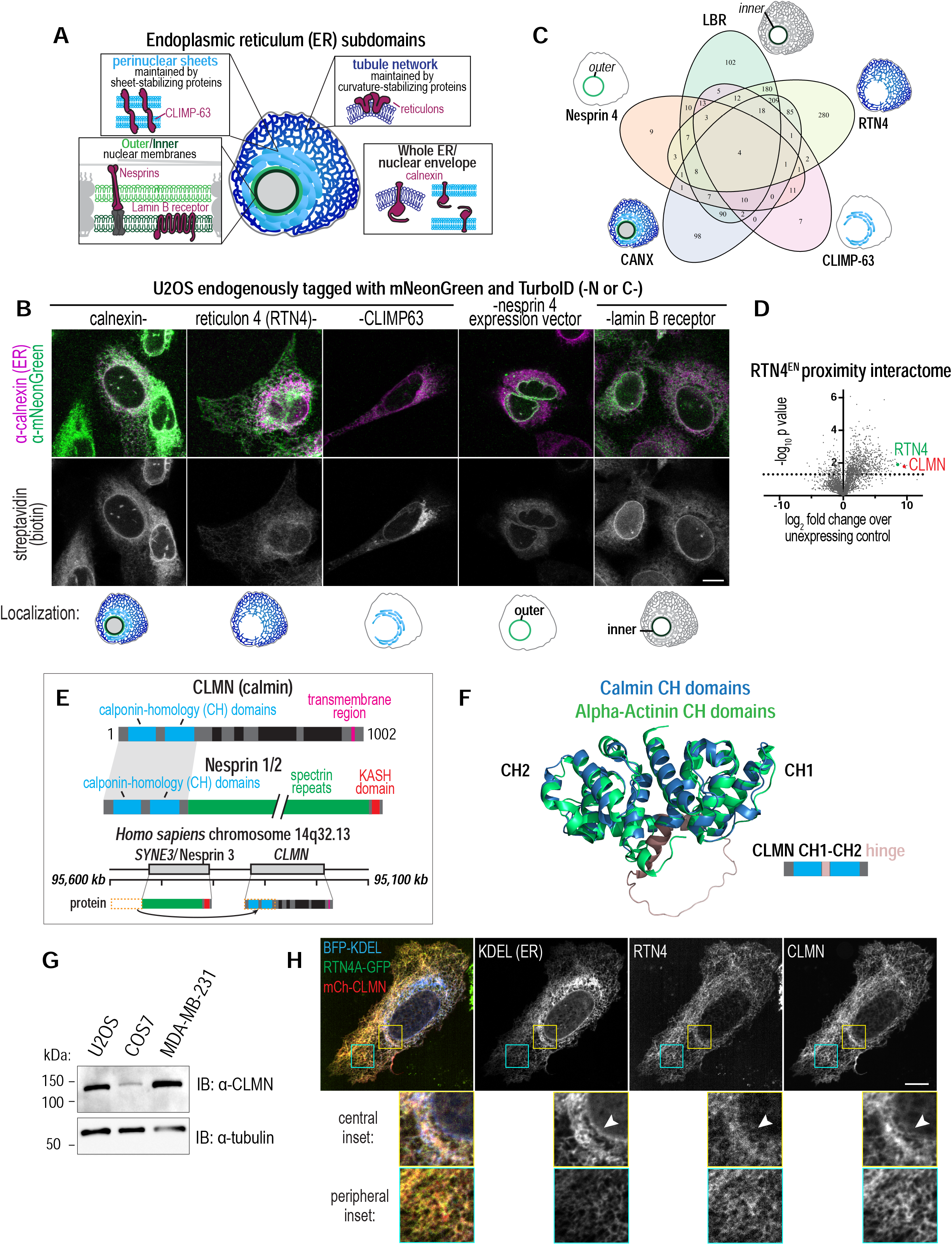
A TurboID-based proteomics screen for ER sub-domain proteins uncovers CLMN as an ER tubule protein. A) Schematic of morphological subdomains of the ER. Indicated proteins were selected as markers for ER sub-domain compartments for proximity labeling proteomics. B) Laser scanning confocal images of U2OS cells treated with 50 µM biotin for 10 minutes prior to fixation and immunolabeling with indicated markers. Scale bar = 10 µm. C) Venn diagram of number of proteins that were identified as positive hits (fold change over un-expressing control > 1 and p < 0.05) in each proximity-labeling proteomics dataset corresponding to ER subdomain markers, and overlap between each list. D) Volcano plot of TurboID-based proteomics showing enriched proteins in the RTN4-TurboID-mNeonGreen endogenously-tagged U2OS cells compared to unmodified U2OS, with log2 fold enrichment on x axis relative to -log10 p values on the y axis. P values, 2-tailed t test of normalized peptide abundances per protein detected (N = 3 experimental repeats). E) Schematic of domain architecture of CLMN relative to Nesprins. Below: schematic of hypothesis of how CLMN arose through a genomic rearrangement with Nesprin 3 (Sequence info from: NC_000014.8 Chromosome 14 Reference GRCh38.p14 Primary Assembly). F) Ribbon structures of calponin-homology domains of CLMN (AlphaFold predicted structure Q6NUQ2) and alpha-actinin (PDB 2EYI^39^) overlaid. Schematic of CLMN CH1CH2/hinge region is shown. G) Immunoblot of indicated markers in whole-cell lysates from indicated cell lines. Representative image from N = 3 experimental repeats. H) Live spinning disk confocal images of U2OS cells transiently expressing indicated markers. Scale bar = 10 µm. Arrows point to area of nuclear envelope and perinuclear ER sheets.

We developed a protocol where cells were exposed to a brief 10-minute pulse of 50 µM biotin^30^, then fixed, lysed, lysates dialyzed to remove excess biotin, and biotinylated proteins enriched by affinity purification on streptavidin beads. Biotinylated proteins were then sent for LC-MS/MS proteomics analysis. Proteomics detected many ER proteins significantly enriched over non-TurboID expressing control cells, and revealed distinct enrichments of specific ER proteins in only some of the TurboID Marker Protein datasets, suggesting they partition into particular ER morphological sub-regions (Figures 1C, S1B-C, Table S1). In line with this, fluorescent staining of the TurboID-expressing Marker Protein cells with fluorescent streptavidin revealed some partitioning of biotinylated proteins into ER subregions corresponding to Marker Proteins, suggesting the biotinylated proteomes reflected the local protein enrichment in different ER morphological subregions (Figure 1B). A dimmer streptavidin signal was present throughout the ER network in some cells, and may represent “blurring” of this subregion enrichment as some biotinylated proteins naturally diffuse across the ER network during the experimental time period.

Among the top hits enriched in the RTN4 TurboID dataset was the protein Calmin (CLMN), a protein of unknown molecular function and subcellular localization (Figure 1D). Comparatively, CLMN was not significantly enriched in the general ER network dataset for CANX nor in the CLIMP-63 sheet proteome, suggesting it partitioned in ER tubules (Figure S1C). CLMN was a minor hit in the LBR dataset, but this may be due to LBR spending part of its lifetime in ER tubules after its synthesis and during its regulated degradation^31,32^ (Figure S1C). CLMN was first identified in mouse embryonic dorsal epidermis as a vertebrate-specific calponin homology (CH) and transmembrane (TM) domain-containing protein localizing to intracellular membranes^22^ (Figure 1E). Further studies confirmed CLMN expression in a variety of tissues, including the mouse testis, liver, kidney, large intestine, and in developing and mature mouse brain in a manner sensitive to all-trans retinoic acid^22,33–36^. Structurally, the CH domains of CLMN are most closely related to those of the LINC complex components Nesprin 1/2^37^ (Figure 1E) and could represent an actin-binding module for CLMN, as CH domains are known to interact directly with actin^38^. Intriguingly, Nesprin 3 lacks CH domains, and its encoding gene *SYNE3* is directly adjacent to *CLMN* in vertebrate genomes, suggesting that the *CLMN* gene arose through a genomic rearrangement event in vertebrates that adopted the CH domains of the ancestor Nesprin 3 gene and recombined it with adjacent sequences to make the new, vertebrate-specific CLMN protein^37^ (Figure 1E). The AlphaFold predicted structure of CLMN supports this hypothesis, in that the CLMN CH1 and CH2 domains closely overlay with the well-characterized CH domains of alpha-actinin, which directly binds to actin^39^ (Figure 1F). The remainder of CLMN before its TM region is predicted to be intrinsically disordered (Figure S2A). We confirmed the broad expression of CLMN by immuno-blot in human U2OS and MDA-MD-231 breast cancer cells as well as African green monkey kidney (COS7) cells (Figure 1G). Thus, CLMN is a vertebrate protein with CH domains that potentially localizes specifically to ER tubules. Its putative actin-binding CH domains, potential ER localization, and large size (1002 amino acids) lead us to hypothesize that CLMN may act as an ER-actin tether, of which none are currently known in animal cells.

### CLMN localizes to ER tubules and F-actin assemblies on the cell basolateral surface

To verify whether CLMN localizes to ER tubules as indicated by the RTN4 TurboID profiling, N-and C-terminally tagged CLMN constructs were generated and transiently expressed in U2OS cells. Live-cell imaging revealed that CLMN colocalized with RTN4-positive tubules but did not appear to decorate the NE or perinuclear ER, consistent with CLMN localizing primarily to curved ER membrane tubules (Figure 1H). Indeed, both N-terminally mCherry tagged and C-terminally GFP tagged CLMN localized to ER tubules and were generally not detectable on the NE (Figure S2B). In addition to its ER tubule localization, we noted that N-and C-terminally tagged CLMN localized to punctae at the basolateral surface of the cell (Figure 2A, Figure S2B, insets). Similarly, mild over-expression of CLMN-GFP also appeared to promote endogenous RTN4, but not CANX, to enrich at these basolateral punctae, supporting the TurboID proteomics that CLMN resides primarily in the RTN4-positive ER tubular network (Figure S3A-B).

**Figure 2:**
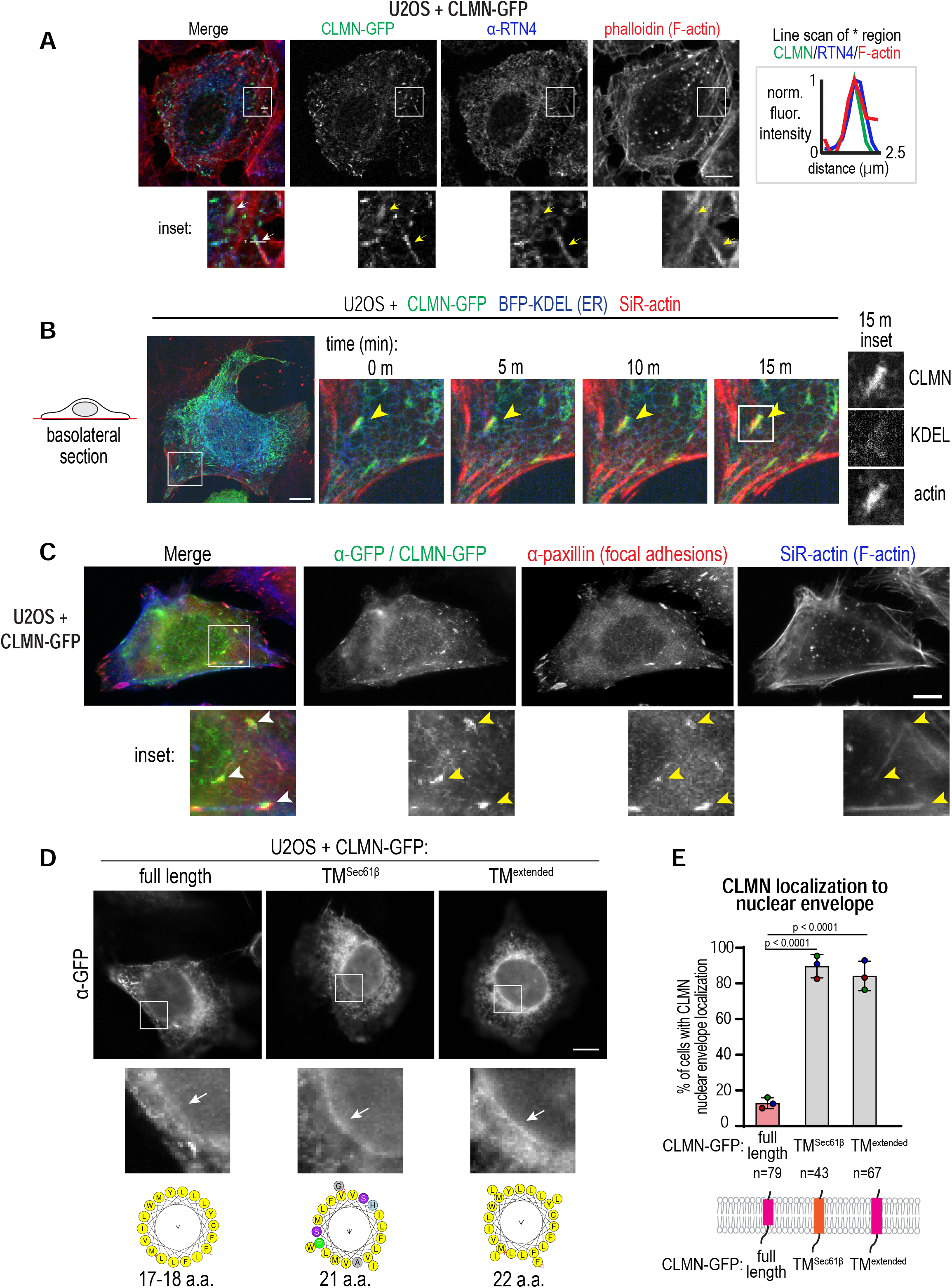
CLMN localizes to ER tubules and basolateral surface punctae and is excluded from the nuclear envelope via its transmembrane domain. A) Laser scanning confocal images of U2OS cells transiently expressing CLMN-GFP treated with DMSO prior to fixation and immunolabeling with indicated markers. Scale bar = 10 µm. Right: Plot of line scan of * region in image across 3 pixel-width line and normalized to minimum/maximum intensities of each marker. B) Live spinning disk time lapse confocal images of U2OS cells transiently expressing or labeled with indicated markers/dyes. Scale bar = 10 µm. Arrows point to stable actin/CLMN puncta. C) Laser scanning confocal images of fixed U2OS transiently expressing CLMN-GFP immunolabeled with indicated markers. Scale bar = 10 µm. Arrows point to CLMN/paxillin-positive foci. D) Laser scanning confocal images of fixed U2OS transiently expressing CLMN-GFP immunolabeled with indicated marker. Scale bar = 10 µm. Arrows point to nuclear envelope area. Below: HeliQuest plots of transmembrane helices from CLMN, Sec61β, and CLMN-TM^extended^. E) Plot of incidences of localization of CLMN to the nuclear envelope in U2OS cells expressing indicated markers as in D. P values, Fisher’s exact tests of pooled incidences. Replicate means ± SDs shown; n = number of cells within N = 3 experimental repeats. Below: schematic of transmembrane domains of transiently-expressed CLMN constructs within a lipid bilayer.

CH domains are known to bind F-actin^40^, so we hypothesized these CLMN-positive basolateral punctae may correspond to F-actin structures at the basolateral cell surface. Indeed, co-staining cells transiently-expressing CLMN with the fluorescent F-actin-binding molecule phalloidin revealed CLMN-GFP punctae frequently colocalized with F-actin-positive structures (Figure 2A) that were approximately 1 µm in width (Figure 2A, line scan). To determine the stability of CLMN basolateral punctae with F-actin, living cells were imaged over a time span of 15 minutes with the live cell imaging-compatible actin-binding dye SiR-actin (Figure 2B, Video S1). CLMN-GFP punctae generally persisted for the duration of the 15-minute time lapse, indicating that CLMN punctae are relatively stable with F-actin (Figure 2B). Thus, CLMN localizes to ER tubules and to F-actin positive structures in the cell periphery.

### CLMN associates with paxillin-positive focal adhesions on the cell surface

Because of the size and stability of these F-actin positive CLMN basolateral punctae, we hypothesized they were associated with focal adhesions (FAs)— F-actin-rich structures on the cell basolateral surface that anchor the cell to its environment^41,42^. Indeed, co-staining of phalloidin and the FA marker paxillin and CLMN revealed CLMN enriched specifically at a subset of paxillin-positive FAs (Figure 2C). Furthermore, CLMN-positive FAs appeared to reside at the ends of F-actin stress fibers, consistent with localization to stress fibers adjacent to cellular adhesions (Figure 2C). Collectively, this supports a model where CLMN associates with the ER tubular network, as well as F-actin and paxillin-positive FAs on the basolateral surface of cells.

### The CLMN transmembrane domain determines its ER tubule localization

Given CLMN’s enrichment at ER tubules and FA structures at the cell surface, we wondered mechanistically how CLMN was enriched in the ER periphery. A feature of proteins that localize to ER morphological subdomains, such as CLIMP-63 to ER sheets or reticulons to tubules, is that they generally do not decorate the NE^8,9,43^. CLMN is not predicted to contain any structural features such as a membrane hairpin that would facilitate enrichment to regions of membrane curvature. However, its TM domain is predicted to be short, with only 17-18 amino acids (Figure 2D). We observed that while it decorated the ER tubule network, it was excluded from the NE (Figure 2D). To determine if the length or composition of CLMN’s transmembrane domain influenced its exclusion from the NE, we swapped it with the TM domain of the ER translocon component Sec61β, which is known to distribute across the entire ER and NE membrane system^9^. Unlike full-length CLMN-GFP, this CLMN-TM^Sec61β^-GFP chimera localized to both the tubular network and the NE in a majority of cells, suggesting the CLMN TM region confers NE exclusion (Figure 2D-2E). As the amino acid composition of Sec61β’s TM domain is not as hydrophobic as that of CLMN, we also generated a CLMN construct with the TM domain extended with 4 more hydrophobic residues (ILLF) to create a 22 amino-acid hydrophobic TM domain. Intriguingly, this TM^extended^ CLMN also now decorated both ER tubules and the NE in the majority of cells (Figure 2D-2E). This suggests that the short length of CLMN’s TM domain is required for CLMN exclusion from the NE.

### CLMN’s CH1-CH2 domains mediate its association with specific pools of F-actin

Next, we dissected how the CH domains and other regions of CLMN influenced its partitioning within the ER periphery and at FAs. The CH domain is an α-helical fold that interacts directly with actin and can impact actin dynamics and cell signaling^40^. In proteins containing type I (CH1) and type II (CH2) CH domains, the CH1 domain directly binds to actin, while the CH2 domain sterically hinders this interaction and thus regulates protein-actin interactions^44–46^. Thus, CH1 and CH2 intramolecular interactions can tune the specificity of the CH1-CH2 domain-containing protein for different F-actin structures within the cell, such as to the front or rear of motile cells^47^. The actin-binding domain (ABD) of Nesprin 2, for example, localizes to the lamellipodium, while the ABD of dystonin localizes specifically to stress fibers^47^.

To determine if CLMN can serve as an ER-actin molecular tether, we conducted structure-function analysis using CLMN truncation mutants (Figure 3A). As expected, full-length CLMN localized to ER tubules and to basolateral punctae in most cells (Figure 3B-3C). Deletion of the CH1 domain or the CH1-CH2 ABD abolished CLMN localization to these basolateral punctae, and instead these truncations localized only to the ER network, suggesting the CH1 and ABD regions are necessary for actin interactions (Figure 3B-3C, ΔCH1, ΔCH1CH2). In contrast, deletion of only the CH2 domain did not impact CLMN’s actin colocalization; CLMN retained ER localization and colocalized with F-actin structures, but was now enriched in actin assemblies away from cell leading edges (Figure 3B-3C, ΔCH2). Intriguingly, this polarization also led to regions of the ER re-localizing to the posterior end of the cell (Figure 3C, ΔCH2, α-RTN4), indicating that CLMN’s actin association may influence how the ER tubular network interacts with actin structures. These data suggest that the CH2 domain of CLMN determines some aspect of specificity in the CLMN-actin interaction and influences where the ER interacts with F-actin. In agreement with previous results^22^, deletion of the CLMN TM segment resulted in it being soluble and cytoplasmically localized (Figure 3B-3C), although traces of F-actin interactions were still observed (Figure 3C, ΔTM). Deletion of the remainder of the C-terminus along with the TM region also resulted in a soluble protein (Figure 3B, ΔC ter). Collectively, this suggests that CLMN’s ABD and its CH1 domain are necessary for F-actin interaction, and its TM region is necessary for ER localization. This supports a model where CLMN functions as an ER-actin tether, with its CH domains governing actin interactions.

**Figure 3:**
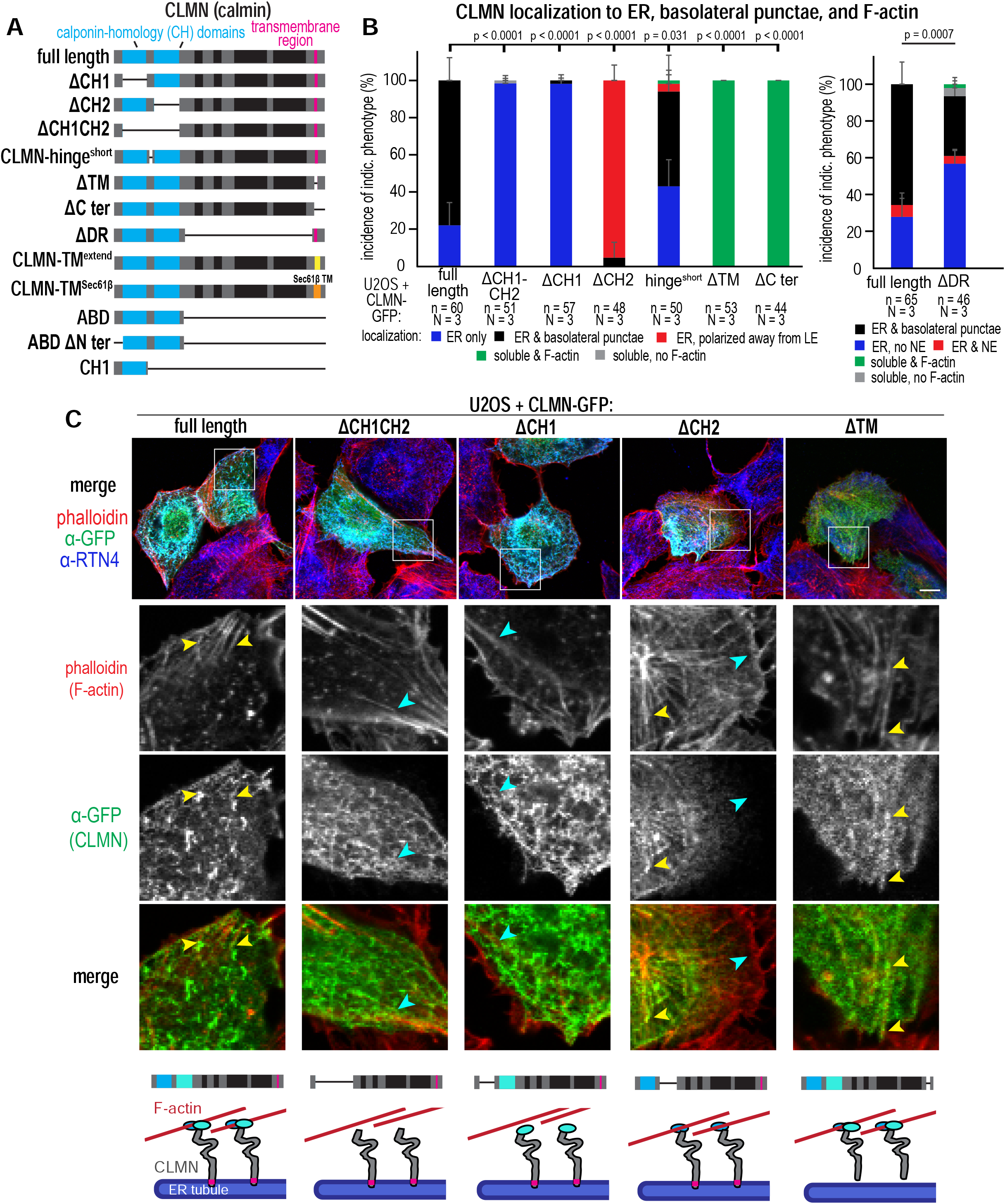
Structure-function analysis reveals that CLMN is an ER-actin molecular tether. A) Schematic of domain architecture of CLMN truncation mutants used to assess structure-function relationships. Black boxes in domain structure indicate predicted disordered regions; yellow box indicates extended transmembrane segment, and orange box indicates Sec61β transmembrane segment. CH = calponin-homology domain; TM = transmembrane domain; C ter = C terminal region; DR = disordered region; ABD = actin-binding domain; N ter = N terminal region. See Methods for exact amino acid changes. B) Plots of incidences of localization of transiently-expressed CLMN to the indicated compartments. n = number of cells within N experimental repeats. Replicate means ± SDs shown. P values = Fisher’s exact test of pooled incidences; each condition compared to full length. C) Laser scanning confocal images of U2OS cells transiently expressing indicated CLMN-GFP vectors prior to fixation and immunolabeling with indicated antibodies/dyes. Yellow arrows indicate areas of CLMN enrichment at F-actin filaments; cyan arrows indicate areas of no/low CLMN enrichment at F-actin filaments. Below: schematic of predicted binding topology based on localization. Scale bar = 10 µm.

Next, we investigated how the CH1-CH2 interactions influenced CLMN interactions with actin. From previous work, the length and composition of the hinge region connecting the CH1 and CH2 domains modulate conformational flexibility between these domains, and ultimately dictate propensity for actin binding^40,45,48^. Of note, structural overlay comparisons between the CH1-CH2 domains of actinin and CLMN suggest CLMN has a significant hinge region connecting these two domains (Figure 1F). To determine whether this hinge region influences CLMN’s interaction with actin structures, we generated a shortened hinge CLMN mutant. Indeed, shortening the CLMN hinge region between the CH1-CH2 domains reduced CLMN localization to basolateral punctae (Figure 3B, hinge^short^; Figure S4A). Since CLMN contains predicted disordered regions (DRs) across the middle of the protein, we also queried whether this was important for proper CLMN subcellular targeting. Deletion of the region encompassing the DRs led to a decrease of CLMN localization to basolateral punctae, possibly supporting a role for this region in assisting in ER-actin tethering and stabilizing actin binding (Figure 3B, ΔDR; Figure S4A).

We also queried whether the ABD of CLMN alone was sufficient for actin interaction *in vivo*, as this region is sufficient for actin binding in other CH1-CH2 proteins^45,47^. Expression of CLMN’s ABD (i.e. CH1+CH2 domains) reveals it as soluble in the cytoplasm, but partially colocalized with F-actin structures, suggesting actin binding (Figure S4B-C). Indeed, co-staining of phalloidin with FA marker paxillin in CLMN ABD-GFP-overexpressing cells revealed the CLMN ABD region localizes to a variety of F-actin structures, including stress fibers, FAs, and F-actin at the leading edge (Figure S4B-C, ABD). In contrast, the ABD of CLMN lacking the CH2 domain (CH1 only) displayed altered actin association, localizing to F-actin structures throughout the cell in a manner biased away from the leading edge and away from stress fibers associated with paxillin, as shown by low overlap between GFP and paxillin signals in a line scan across the cell periphery (Figure S4B-S4C, CH1). We also examined how loss of the TM region influences CLMN localization. Full-length CLMN lacking the TM domain phenocopies the localization of the ABD alone, but appeared more weakly co-localized with F-actin, suggesting that the regions of CLMN outside of the ABD may also serve to constrain or fine-tune F-actin binding (Figure S4C, ΔTM). Finally, since the N-terminus upstream of the CH1 domain has also shown to support actin binding in other proteins, we investigated its importance in CLMN binding to actin structures^45,49^. Indeed, deletion of the N-terminus upstream of CLMN’s ABD led to reduced F-actin localization, in agreement with previous studies, suggesting this region contributes to actin interactions (Figure S4D). Altogether, this is consistent with the role of CLMN as a molecular tether connecting the ER network and actin structures, as protein tethers are operationally defined as large (i.e. hundreds of amino acids) proteins with clear anchoring or TM regions connecting two or more distinct cellular structures^50,51^. It also supports a model where the CH1 and CH2 domains, like in other actin-binding proteins, may function to fine-tune CLMN’s interactions with actin structures to provide specificity for CLMN’s interactions between the ER and actin network.

### CLMN is required for normal F-actin cell morphology and cell migration

To determine the effects that CLMN-mediated tethering of the ER and F-actin may have on F-actin organization, we treated U2OS cells with 30nM CLMN siRNA for 48 h and co-stained for F-actin (with phalloidin) and ER tubules (RTN4) (Figure 4A). We confirmed siRNA-mediated CLMN knockdown by qPCR and Western blot (Figure S5A-S5B). Strikingly, CLMN-depleted cells exhibited significant changes in F-actin structures compared to control cells. CLMN-depleted cells exhibited accumulations of peripheral F-actin bundles near the cell periphery, likely corresponding to bundled actin stress fibers (Figure 4A-4B).

**Figure 4:**
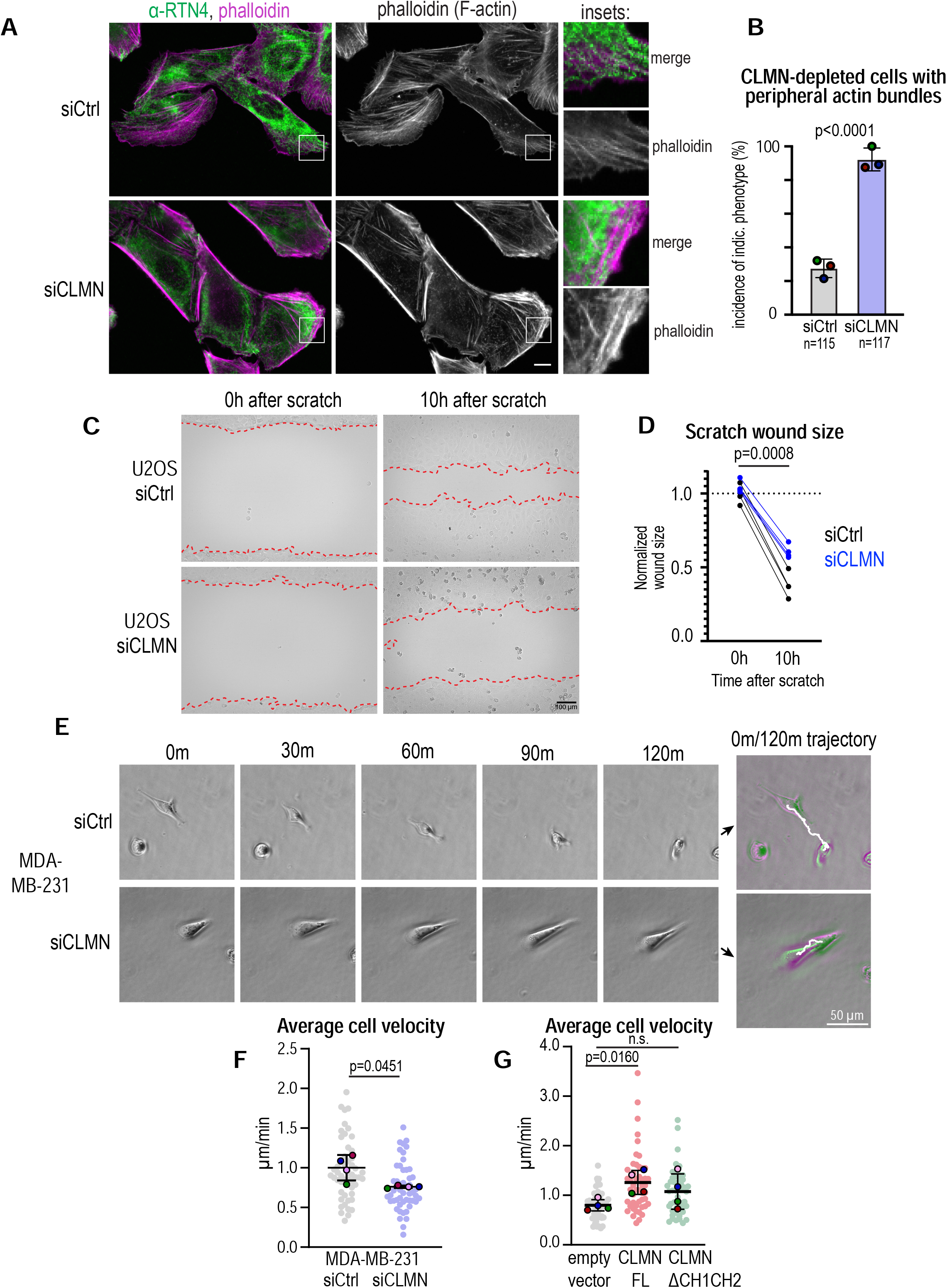
CLMN depletion results in actin cytoskeletal reorganization and reduced cell motility. A) Laser scanning confocal images of U2OS cells treated with indicated siRNAs prior to fixation and immunolabeling with indicated antibodies/dyes. Scale bar = 10 µm. B) Plot of incidences of indicated phenotypes in U2OS cells as in A. P value, Fisher’s exact test of pooled incidences. N = number of cells within N = 3 experimental repeats. Replicate means ± SDs shown. C) Phase contrast images of live U2OS cells treated with the indicated siRNA and mitomycin to inhibit proliferation, then subjected to scratch assay. Scale bar = 100 µm. Red dotted lines indicate wound edges. D) Plots of wound size over time in cells treated as in C, normalized to control average size at t = 0 hours. P value = paired t test of replicate means of 0 h wound areas – 10 h wound areas. Replicate means shown; n = 2 fields of view each per N = 4 experimental repeats. E) Phase contrast time lapse images of MDA-MB-231 cells treated with the indicated siRNAs and allowed to freely migrate in low-density conditions. Right, composite overlays of first and last time points with frame-by-frame trajectories shown in white. Scale bar = 50 µm. F) Plots of average velocities of cells treated as in F. Individual cell velocities (filled circles) and replicate means (outlined circles) ± SDs from N = 4 experimental repeats shown. P value = paired t test of replicate means. G) Plots of average velocities of cells transiently expressing indicated vectors. Individual cell velocities (filled circles) and replicate means (outlined circles) ± SDs from N = 4 experimental repeats shown. P values = paired t tests of replicate means.

Since this actin reorganization would potentially impact cell motility, we next conducted cell migration assays. We first conducted a scratch assay with control and CLMN-depleted U2OS cells to determine the impact of CLMN expression on directional cell migration (Figure 4C). After 10 h post-scratch induction, CLMN-depleted cells made significantly less progress in migrating toward the wound bed compared to control cells, suggesting that loss of CLMN impacted directional cell movement (Figure 4D).

To determine if CLMN impacts migration in cell lines other than U2OS, the highly migratory breast cancer cell line MDA-MB-231 was used to track the un-constrained migration of single cells over time^52^ (Figures 4E, S5B, Video S2). Strikingly, over 2 h MDA-MB-231 cells depleted of CLMN exhibited a reduced average velocity compared to control cells (Figure 4F). We also examined how CLMN over-expression influenced cell motility. Intriguingly, overexpression of full-length CLMN, but not CLMN lacking the CH1-CH2 ABD, resulted in an increased average velocity compared to cells expressing a control vector (Figure 4G). Taken together, these data support a role for CLMN in enabling efficient cell migration and promoting F-actin dynamics.

### CLMN localizes to a sub-population of FAs, and its depletion alters FA density on the basolateral surface

An important cellular component for cell motility are cellular adhesions, which act as multiprotein complexes that link actin stress fibers to the extracellular matrix^53^. Cellular adhesions consist of layers of proteins including integrins interacting with the extracellular matrix, as well as intracellular signaling and force transduction layers that mediate actin interactions^41^. Since CLMN colocalized with paxillin-positive cellular adhesions, and its depletion altered cell movement, we predicted that CLMN influenced some aspect of adhesion homeostasis. To test this, we first examined whether CLMN localized to all cellular adhesions, or to specific subsets. We co-stained for paxillin in cells expressing CLMN-GFP. Surprisingly, CLMN did not co-localize with all paxillin foci. CLMN did associate with some adhesions at the cell peripheral edges, but appeared most enriched at paxillin foci generally underneath the nucleus and cell body (sometimes denoted as ventral adhesions^54^) (Figure 2C). During mesenchymal cell migration, adhesion dynamics are polarized to adhesions formed at the cell leading edge, and they disassemble shortly thereafter or mature into FAs. In this process, more mature adhesions are disassembled underneath the cell body or at the trailing edge of the cell^54^. Because CLMN is enriched at ventral adhesions closer to the cell mid-body and trailing edge, we reasoned CLMN may be biased toward association with disassembling adhesions.

Given the strong effect of CLMN depletion on cell migration in combination with the selective colocalization with mature FAs, we hypothesized that CLMN may play a role in adhesion disassembly. To examine this, we assessed whether CLMN depletion altered the size or surface density of paxillin-positive foci. Cells migrating toward a wound were co-stained for paxillin and phalloidin, and paxillin signal was segmented using Weka machine learning segmentation^55^ (Figure 5A). Indeed, CLMN-depleted cells exhibited an increased number of paxillin foci per cell area compared to siCtrl cells (Figure 5B). Importantly, this change in adhesion density could be suppressed by transiently expressing full-length CLMN-GFP, but not CLMN-GFP lacking any component of the ABD or lacking the transmembrane domain, indicating that CLMN ER-actin tethering was necessary for function (Figure 5C). CLMN depletion or re-expression did not influence paxillin foci size (Figures 5D-E). Collectively, these data suggest that the ER-actin tethering function of CLMN plays a role in adhesion dynamics, and that CLMN loss impacts some aspect of paxillin-positive cellular adhesion dynamics.

**Figure 5:**
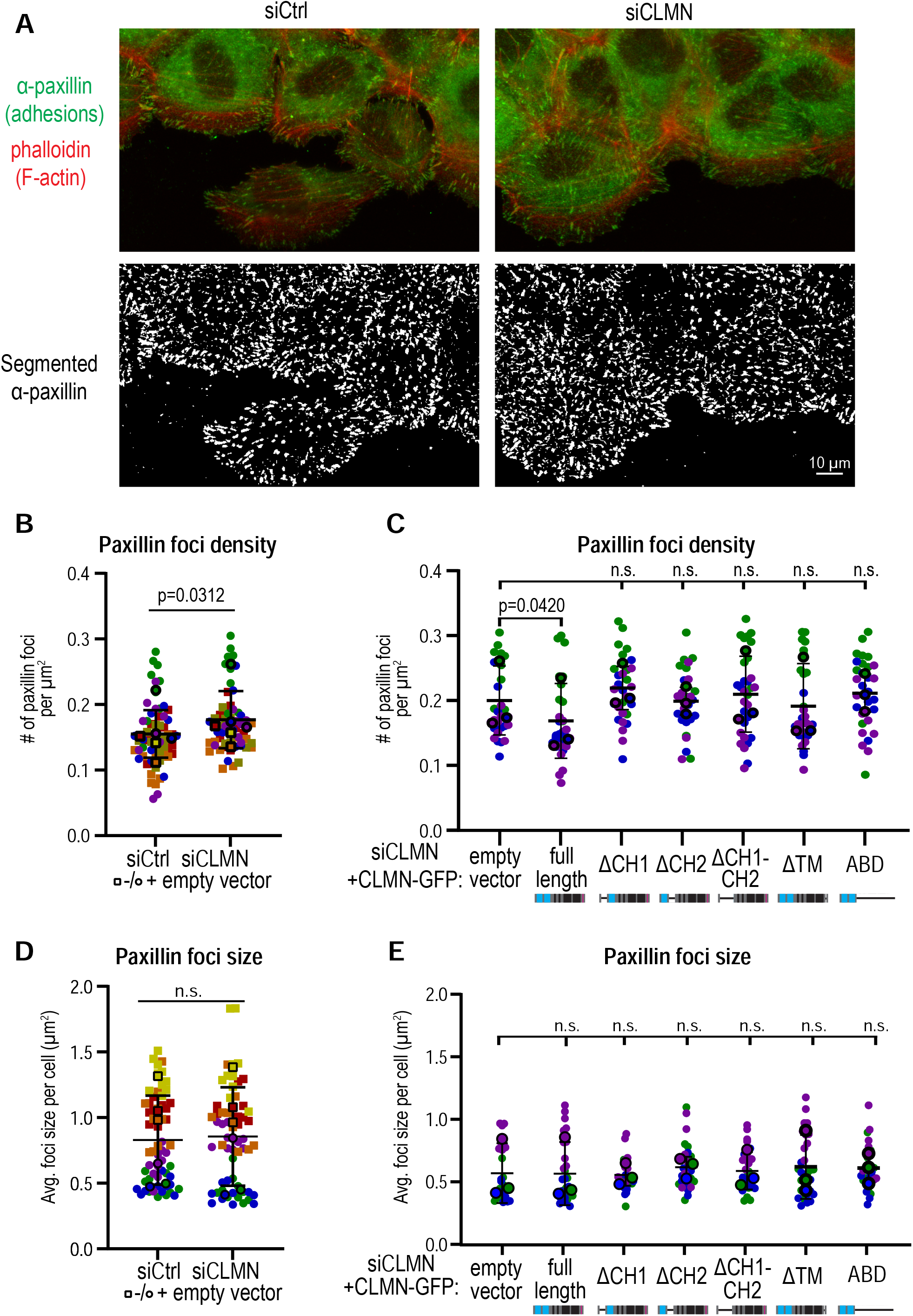
CLMN expression controls cellular adhesion density. A) Above: Laser scanning confocal images of cells treated with the indicated siRNAs and allowed to migrate to a scratch wound for 2 hours before fixation and immunolabeling with the indicated antibodies/dyes. Below: Weka segmentation of paxillin signal in the above images to detect cellular adhesions. Scale bar = 10 µm. B) Plot of number of paxillin foci per cell area in cells treated as in A. Individual cell paxillin foci densities (filled shapes) and replicate means ± SDs (outlined shapes) from N = 6 experimental repeats shown. Square points versus circle points indicates whether or not cells were expressing empty vector within that experimental repeat. P value = Wilcoxon matched-pairs signed rank test of replicate means. C) Plot of number of paxillin foci per cell area in cells treated with indicated siRNA and transiently expressing indicated vectors. Individual cell paxillin foci densities (filled shapes) and replicate means (outlined shapes) ± SDs from N = 3 experimental repeats shown. Note that siCLMN expressing empty vector data also appears in 5B. P values = repeated measures ANOVA of pre-defined pairs with Geisser-Greenhouse correction for non-sphericity and Šídák correction for repeated measures. D) Plot of average paxillin foci size in cells treated with indicated siRNAs. Individual cell values (filled shapes) and replicate means ± SDs (outlined shapes) from N = 6 experimental repeats shown. Square points versus circle points indicates whether or not cells were expressing empty vector within that experimental repeat. P value = paired t test of replicate means. E) Plot of average paxillin foci size in cells treated with indicated siRNA and transiently expressing indicated vectors. Individual cell values (filled shapes) and replicate means (outlined shapes) ± SDs from N = 3 experimental repeats shown. Note that siCLMN expressing empty vector data also appears in 5D. P values = repeated measures ANOVA of pre-defined pairs with Geisser-Greenhouse correction for non-sphericity and Šídák correction for repeated measures.

### ER-actin CLMN tethering can drive ER association with peripheral actin structures

Given that CLMN appeared to mediate ER tubule-actin tethering, and CLMN depletion led to the accumulation of paxillin-positive cellular adhesions, we hypothesized that CLMN mediated the recruitment of ER tubules to adhesions to influence their dynamics, and ultimately cell migration. To begin to address this, we interrogated how CLMN influenced the association of ER tubules with highly dynamic adhesions. We conducted live-cell imaging in the cell periphery (close to the leading edge) where the ER network is virtually absent and provides the greatest resolution for imaging ER-actin dynamics. We imaged living cells transiently expressing either an empty mCherry vector, or over-expressing mCherry-CLMN together with the ER marker BFP-KDEL. Strikingly, mCherry-CLMN over-expressing cells exhibited enrichments of ER tubules in regions of the cell periphery rich in peripheral F-actin filaments (Figures 6A, S6A). Of note, these peripheral ER-actin associations were rare in control (mCherry vector) cells. Quantification revealed that in cells expressing mCherry-CLMN, the proportion of cells with ER tubule localization into the peripheral cell edges increased by more than two-fold (Figure 6B). This suggests that CLMN over-expression was sufficient for ER tubule localization to the cell periphery that is rich in dynamic actin structures.

**Figure 6:**
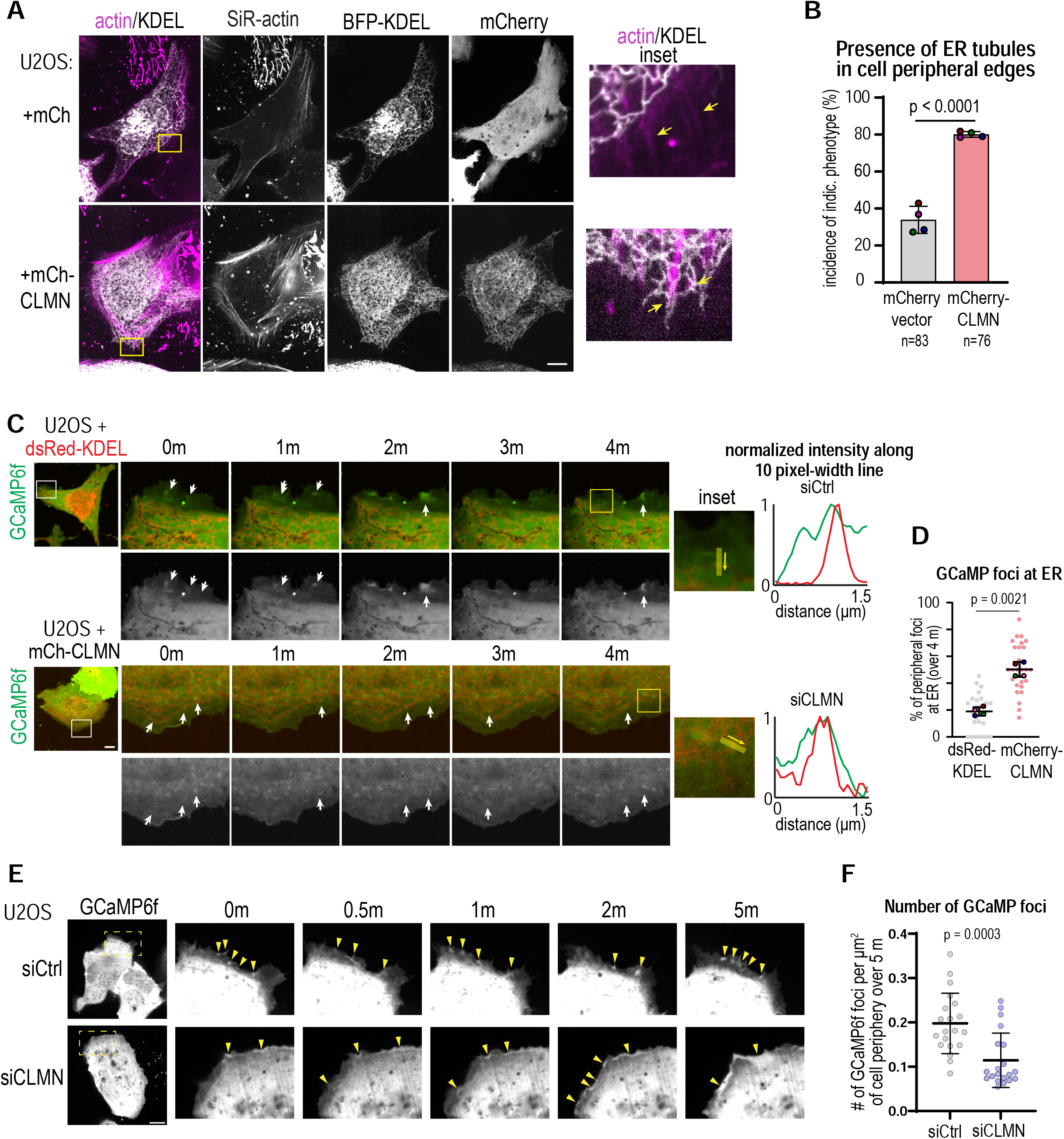
CLMN expression controls ER targeting and influences cellular calcium dynamics. A) Live spinning disk confocal images of U2OS cells transiently expressing the indicated vectors and labeled with SiR-actin. Scale bar = 10 µm. Inset arrows show peripheral actin fibers that are or are not connected to ER tubules. B) Plot of quantification of incidence of ER tubules extending into cell peripheral edges of cells defined by drop-off of cytosolic GCaMP signal in cells expressing indicated vectors (see Methods). n = number of cells within N = 4 experimental repeats; replicate means ± SDs shown. P value = Fisher’s exact test of pooled incidences. C) Live spinning disk confocal time lapse images of U2OS cells transiently expressing the indicated vectors. Scale bar = 10 µm. Arrows point to GcaMP6f foci. Right: line profiles of fluorescent intensity normalized to minima/maxima. D) Plot of percentage of GCaMP foci overlapping with ER signal over the time lapse. Values from individual cells (gray circles) and replicate means ± SDs (colored circles) from N = 4 experimental repeats shown. P value = paired t test of replicate means. E) Live spinning disk confocal time lapse images of cells treated with indicated siRNAs and transiently expressing indicated marker. Scale bar = 10 µm. Arrows point to GCaMP6f foci. F) Plot of number of GCaMP6f flickers per cell area in cells treated as indicated. Values from individual cells (gray circles) from N = 4 experimental replicates and pooled means ± SDs are shown. P value = Mann-Whitney test.

### CLMN ER tethering influences intracellular calcium dynamics in the cell periphery

Next, we investigated how CLMN influences ER-actin functional crosstalk. The ER is a major reservoir of intracellular calcium that serves to amplify and maintain homeostasis of intracellular Ca^2+^ for signaling^56^. Intracellular Ca^2+^ plays a major role in adhesion disassembly and cell migration^57,58^. Increased local Ca^2+^ results in increased residency of focal adhesion kinase (FAK) at adhesions, which will drive adhesion disassembly via activation of the focal adhesion protease calpain, as well as activation of effectors like IQSec1 and the GTPase Arf5 that mediate adhesion disassembly^59–61^. Store-operated calcium entry (SOCE) at ER-plasma membrane contacts mediated by STIM1 and Orai1^62–64^ follows ER calcium store depletion and is targeted to disassembling focal adhesions^61,65^. However, so far it has been unclear which roles ER-stored calcium may play at adhesions and how the ER is targeted to adhesions. Based on our observations, we hypothesized that CLMN tethering of the ER to actin structures adjacent to cell may facilitate targeted Ca^2+^ signaling at adhesions to promote disassembly.

To monitor the influence of CLMN on local Ca^2+^dynamics, we used a sensitive genetically encoded Ca^2+^ biosensor with fast kinetics, GCaMP6f^66^, to assess local changes in Ca^2+^ at the cell periphery over time (Figure 6C). However, changes in fluorescence at the cell periphery could also be attributed to plasma membrane invaginations and other topological changes, so as a control we co-labeled cells with the plasma membrane stain CellMask and verified that GCaMP6f local increases in fluorescence did not overlap with plasma membrane marker local fluorescence increases (Figure S6B).

Next, we examined how CLMN impacted local GCaMP6f Ca^2+^ burst events (denoted as foci here). Strikingly, we observed that transient expression of mCh-CLMN, but not the ER luminal marker dsRed-KDEL, led to an increase in the incidence of GCaMP6f foci at the cell periphery that closely associated with the ER (Figure 6C-D, S6C). This suggests that CLMN over-expression is sufficient to alter Ca^2+^ burst dynamics in the cell periphery. Given that CLMN over-expression also leads to increased ER tubules at the cell periphery, we speculate that increasing ER occupancy in a given cell area leads to increased Ca^2+^ signaling associated with the ER there.

Next, we examined how CLMN depletion altered Ca^2+^ signaling events in the cell. We again imaged GCaMP6f in cells siRNA-depleted of CLMN (Figure 6E, Video S3). Strikingly, we observed that CLMN-depleted cells now exhibited fewer GCaMP6f burst events at the cell periphery over the 5-minute time course compared to control cells (Figure 6F). This suggests that CLMN is necessary for maintaining regular Ca^2+^ signaling near adhesions, and that its depletion alters local Ca^2+^ dynamics. Collectively, it supports the model where CLMN facilitates ER-actin contact near focal adhesions to support local Ca^2+^ signaling, and ultimately proper adhesion disassembly in cell migration.

## Discussion

Organelle-based protein compartmentalization is a fundamental organizational principle of Eukaryotes. Recent advances in imaging and biochemical techniques reveal that proteins and complexes can partition into morphologically unique subdomains within organelles. While some of these, such as mitochondrial cristae are well-described, how ER proteins are organized within different morphological regions of the ER network remains unclear. Using TurboID-based biochemical proteomics screening, we have mapped the local protein landscape of ER and NE subdomains in human U2OS cells. Our proteomics detected the uncharacterized protein CLMN as particularly enriched in RTN4-positive ER tubules. We found that CLMN acts as an ER tubule-actin tether and show that ER-actin tethering is required for proper F-actin architecture, focal adhesion dynamics, and, ultimately, cell migration (Figure 7, above). This is the first, to our knowledge, description of the ER peripheral network being involved in F-actin dynamics and cell motility.

**Figure 7:**
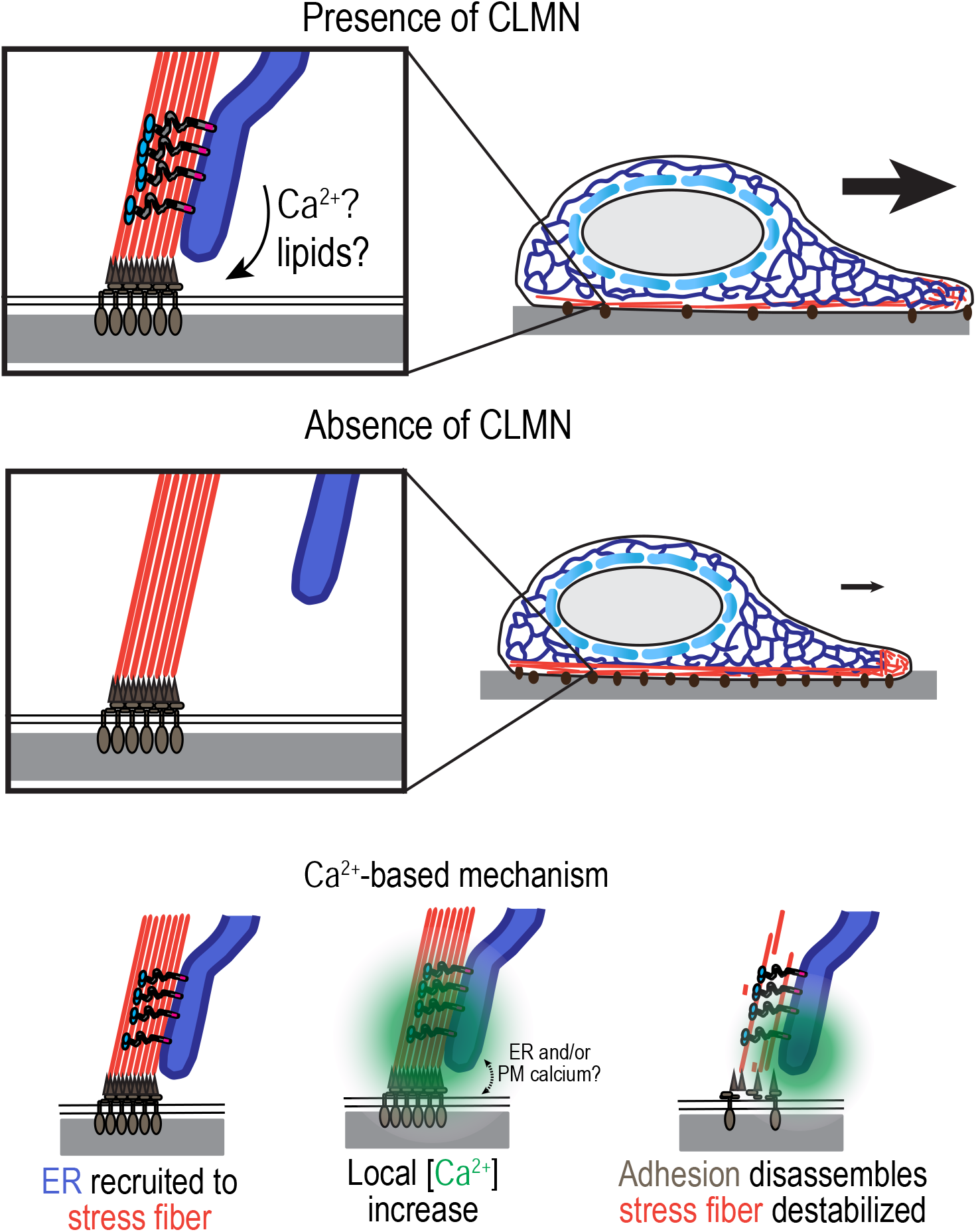
Proposed mechanism for how ER-actin tethering by CLMN facilitates cell migration. Schematic of working model and proposed mechanism for how ER-actin tethering by CLMN controls stress fiber and cellular adhesion dynamics to facilitate proper cell migration. Below: proposed mechanism for how ER-actin tethering by CLMN could facilitate local Ca^2+^ signaling (sourced from the ER or extracellular space) supporting adhesion disassembly.

Cell adhesions and motility are closely regulated by intracellular Ca^2+^ concentration ^57,61,65^. Here, we show that CLMN loss perturbs local Ca^2+^ bursts in the cell periphery, as well as adhesion disassembly and F-actin architecture. We propose that ER-actin tethering spatiotemporally regulates local Ca^2+^ signaling at adhesions to promote disassembly (Figure 7, below). In one possible model, the ER is recruited to adhesions destined to disassemble, and an increase in local Ca^2+^ released from the ER promotes downstream processes that lead to adhesion disassembly and stress fiber destabilization (Figure 7, below). Of note, a canonical mechanism for adhesion disassembly posits that local Ca^2+^ is supplied from the cell surface through Orai1-dependent Ca^2+^ influx^61,65^. Our work presents an alternative non-canonical model where ER tubules can be recruited directly to adhesions via CLMN tethering to facilitate local Ca^2+^ signaling. In this model, CLMN loss would lead to focal adhesion and stress fiber accumulation, and migration would be impaired—all of which are phenotypes observed upon CLMN depletion. Future studies will be needed to further dissect how ER recruitment to FAs influences Ca^2+^ signaling for FA dynamics.

It is also possible that ER-actin tethering by CLMN directly affects actin dynamics in stress fibers or other F-actin structures. For example, it is known that ER-associated actin is required for mitochondrial fission and fusion^67–69^. The ER-associated formin INF2 performs actin polymerization at sites of mitochondrial fission and fusion that is thought to provide mechanical support for membrane dynamics^68^. CLMN may play a role by tethering F-actin filaments to the ER to facilitate their activity at sites of mitochondrial fission and fusion. Further, CLMN could assist in the wrapping of ER around ventral stress fibers during migration-associated nuclear positioning^70^. Another possible role for ER-actin tethering by CLMN is maintaining ER fine architecture. In mitosis, for example, connections between the ER and actin play a dominant role in determining the ultrastructure of the ER^71^, and actin also recruits ER tubules to the reassembling nuclear envelope, although it is not known how these ER-actin connections are achieved^72^. In interphase, mild short-term depolymerization of actin with Latrunculin A leads to a depletion of small peripheral ER sheets, which is dependent on myosin 1c-based ER transport^73^. Our data show that increasing the extent of ER-actin tethering by CLMN overexpression leads to increased ER innervation of peripheral actin fibers. Thus, these data support a role for ER-actin connections in influencing ER structure, which may be mediated by CLMN.

CLMN was identified as being comparatively enriched in proteomics dataset associated with RTN4 (ER tubules or curved membranes). It is unclear how CLMN is enriched to ER tubules specifically, although our findings indicate that its TM domain length confers exclusion from the nuclear envelope. It remains to be seen whether CLMN interacts directly with RTN4 or other ER tubule proteins to lead it to be enriched at curved ER membranes.

CLMN has previously been identified as a protein that is increased in expression in neuroblastoma cells upon all-trans retinoic acid treatment, which induces differentiation into neuronal-like cells^34–36^. CLMN upregulation in this context is thought to promote cell cycle exit^36^, although the mechanism for how this would occur is unclear given its molecular function as an ER-actin tether. It is possible that CLMN performs multiple functions in different cell types to enable cell differentiation. In mice, CLMN is expressed in a variety of tissue types including testis, liver, kidney, large intestine, and brain^22,33–36;^ the function of CLMN in the context of a variety of tissues remains to be understood.

In this study, we have identified the first ER-actin molecular tether, and shown it influences adhesion disassembly and cell motility. This work informs two critical questions within our understanding of organelle biology: first, how division-of-labor occurs between ER structural sub-domains to affect cytoskeletal dynamics, and how the ER-actin interaction promotes a spatially selective interface between ER and FAs that is implicated in the regulation of cellular motility. As maintenance of ER tubules is critical for development of neuronal, muscle, and other tissues, and cell migration underlies proper development, these findings have the potential to inform multiple areas of our understanding of the cell biology of human tissue development and maintenance.

### Limitations of the study

Our understanding of how ER recruitment influences Ca^2+^ and cytoskeletal dynamics at FAs underneath the cell body (as opposed to at the leading edge) is limited by the challenge of spatiotemporally resolving specific ER recruitment events to these adhesions. Similarly, we are limited in our spatial understanding of whether Ca^2+^ is sourced from the ER and/or extracellular space. Thus, we are currently unable to make a direct conclusion about how CLMN localized specifically at basolateral punctae under the perinuclear region plays a role in Ca^2+^ dynamics and adhesion disassembly. Additionally, this study focused on defining the roles of CLMN in adhesion and actin dynamics in cultured human cells. CLMN appears to be broadly expressed, but its roles in different primary cells such as neurons or in a whole-organism context requires further investigation. Future studies will determine how CLMN expression affects cell migration in the context of 3-dimensional migration or within the context of a whole-organism or multiple tissue types.

## Author Contributions

H.M.: Conceptualization, Formal analysis, Investigation, Writing – Original Draft, Visualization; T.I.: Conceptualization, Writing – Original Draft; B.P.: Visualization, Writing – Original Draft; G.D.: Conceptualization, Resources, Writing – Original Draft; W.M.H.: Conceptualization, Resources, Writing – Original Draft, Visualization, Supervision, Project administration, Funding acquisition.

## Acknowledgements

We thank J. Friedman, W. Prinz, J. Liou, and M. Mettlen (UT Southwestern) for critical discussion guiding aspects of this study. We thank E. Reynolds, M. Marlar-Pavey, M. Mettlen, K. Luby-Phelps (UT Southwestern Quantitative Light Microscopy Core Facility), and A. Lemoff (UT Southwestern Proteomics Core Facility) (UT Southwestern) for assistance with experimental design and/or for core facility use. This work was supported by NIH NIDDK grant DK126887, NIGMS grant GM119768, the Welch Foundation I-1873, and the UT Southwestern Medical Center Endowed Scholars Program to W.M.H. and by NIH NIGMS GM145399 to G.D. Additional support is by NIH (T32 DK007307) supporting H.M. and by NIH 1S10OD028630-01 to Katherine Luby-Phelps for the UT Southwestern Quantitative Light Microscopy Core Facility.

---

**Supplementary Figure 1.**
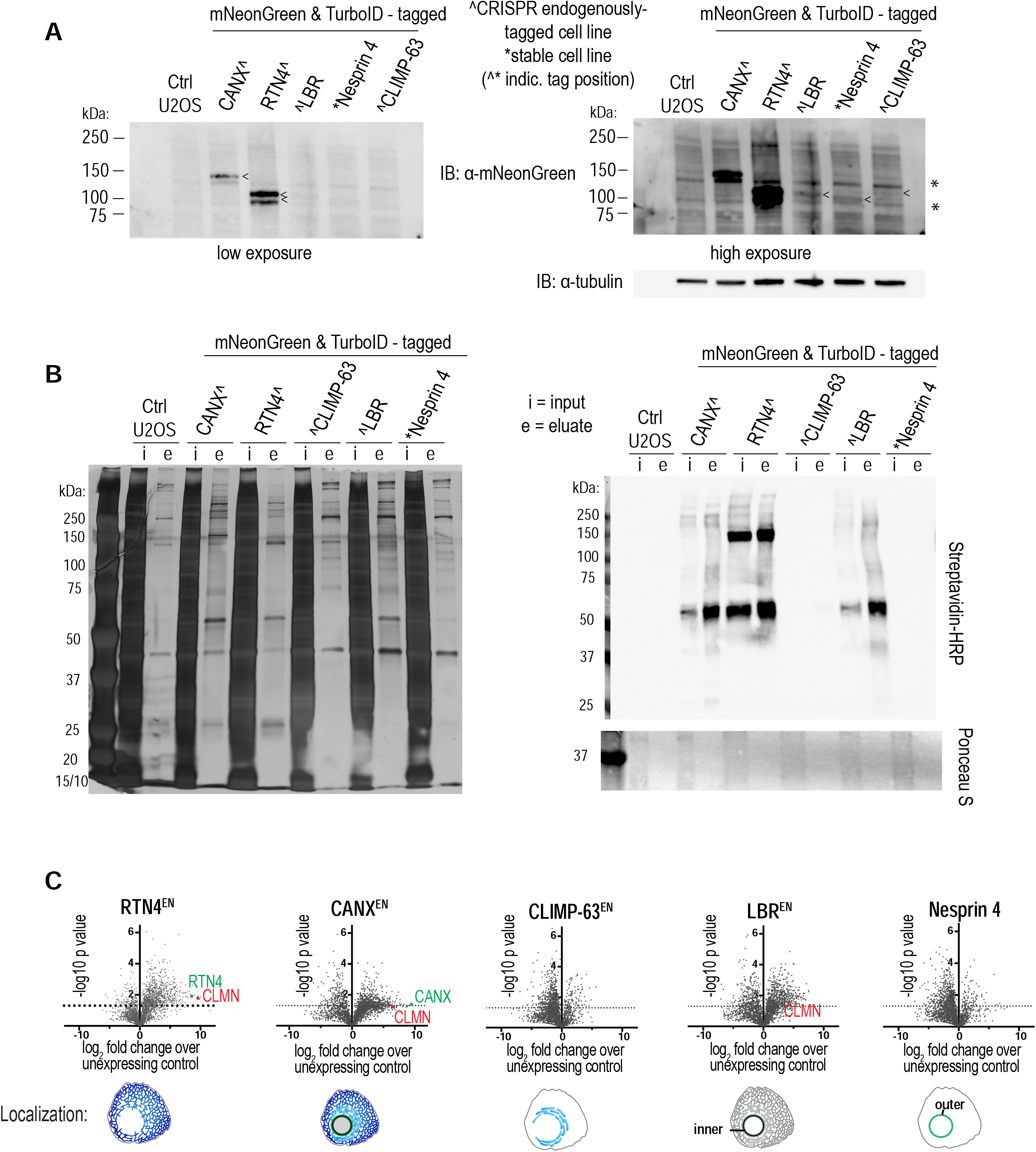
associated with Figure 1: Proteomic analysis of ER sub-domains using TurboID-based proximity labeling. A) Immunoblots of whole-cell lysates from indicated cell lines using indicated antibodies. Arrowheads point to specific bands within lanes; asterisks indicate non-specific bands occurring in every condition. Note pixel saturation of bands in high exposure blot in CANX^ and RTN4^ lanes. Approximate expected molecular weights based on Expasy sequence calculation (kDa): CANX 131, RTN4 104/106, LBR 134, Nesprin 4 107, CLIMP-63 129. LBR tagging was additionally confirmed by genomic DNA PCR due to anticipated size difference (see Methods). B) Silver stain (left) and streptavidin-HRP immunoblot (right) of un-enriched inputs and streptavidin-enriched eluates (biotinylated proteins) in samples from indicated cell lines that were submitted for proteomic analysis. C) Volcano plots of TurboID-based proteomics showing enriched proteins in the indicated U2OS cell lines compared to unmodified U2OS, with log2 fold enrichment on x axis relative to -log10 p values on the y axis. P values, 2-tailed t test of normalized peptide abundances per protein detected (N = 3 experimental repeats).

**Supplementary Figure 2.**
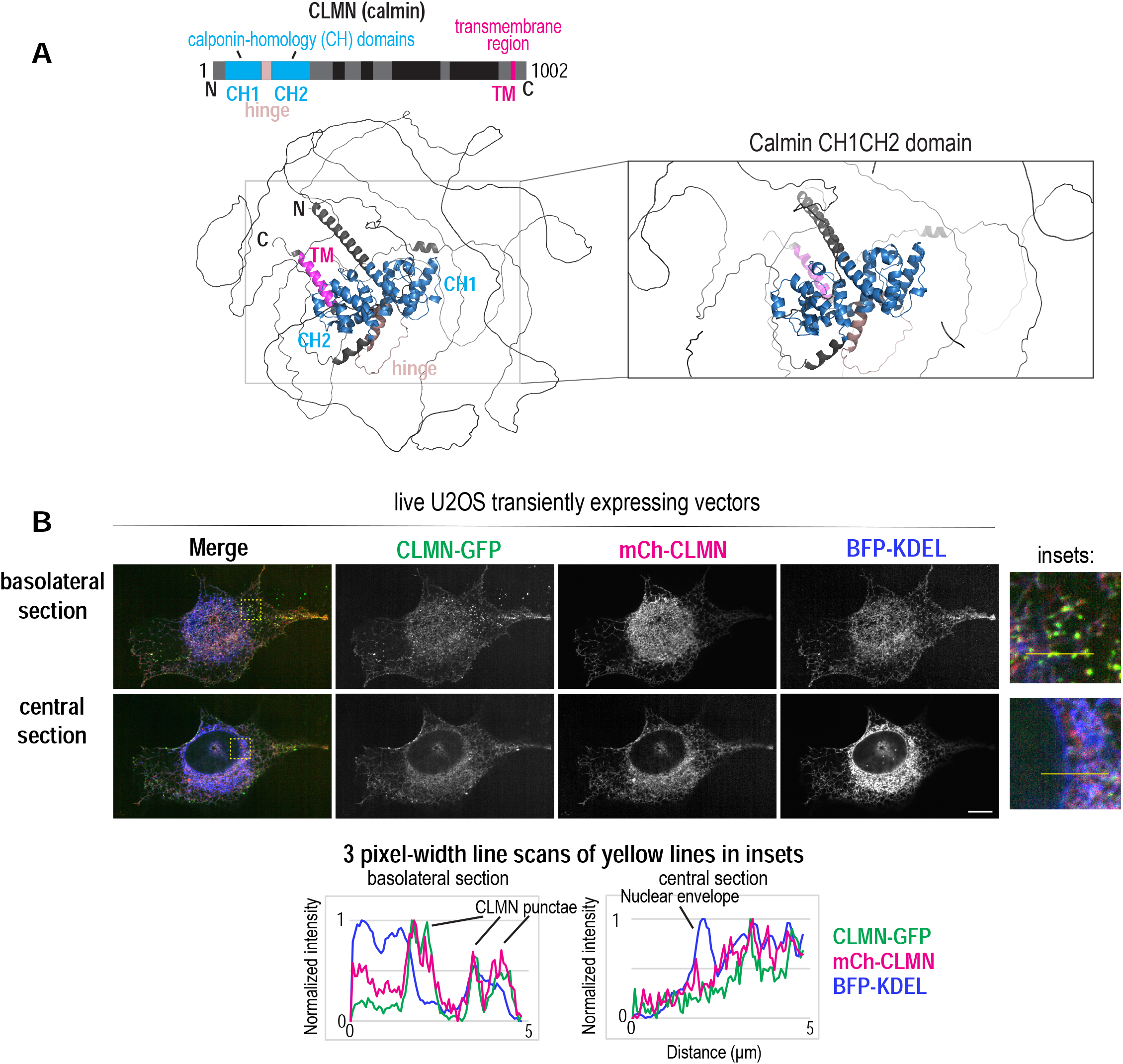
associated with Figure 1: Predicted structure of CLMN and localization of N-terminally and C-terminally tagged CLMN. A) Ribbon structures of AlphaFold predicted structure of CLMN (predicted structure Q6NUQ2) colored by domain as in above schematic. B) Live spinning disk confocal images of U2OS cells transiently expressing indicated vectors. Scale bar = 10 µm. Below: 3 pixel-width line scans along yellow lines indicated in insets. Fluorescent intensities are normalized to minima/maxima.

**Supplementary Figure 3.**
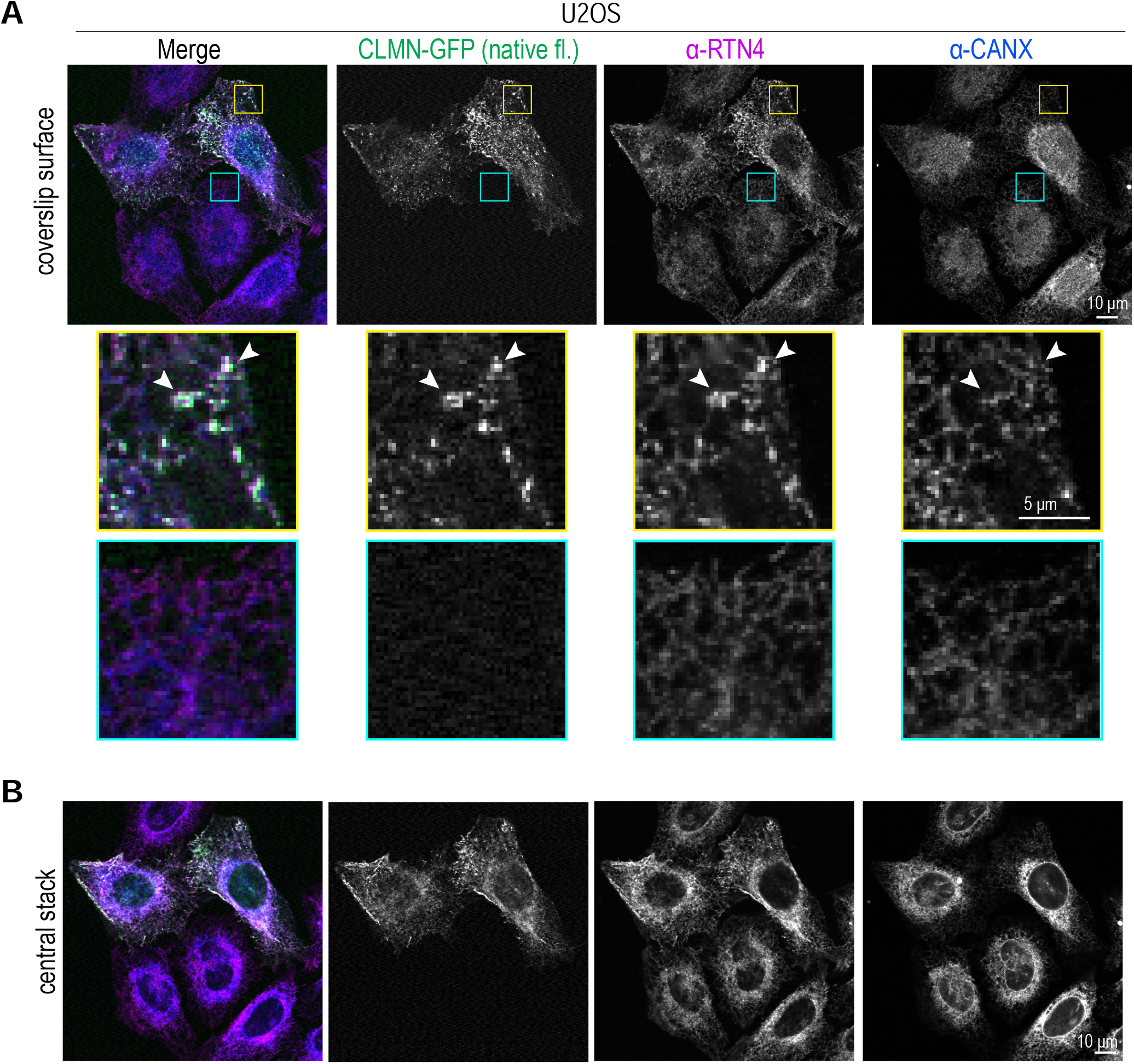
associated with Figure 2: RTN4 localization to CLMN punctae. A-B) Laser scanning confocal images of U2OS cells transiently expressing indicated marker and fixed and immunolabeled with indicated antibodies. Basolateral (coverslip-level) section and central section shown. Scale bars = 10 µm for large images and 5 µm for insets. Arrows indicate CLMN basolateral foci. Yellow boxes indicate areas with CLMN punctae in CLMN-overexpressing cell, while cyan boxes indicate analogous areas in non-CLMN-overexpressing cells.

**Supplementary Figure 4.**
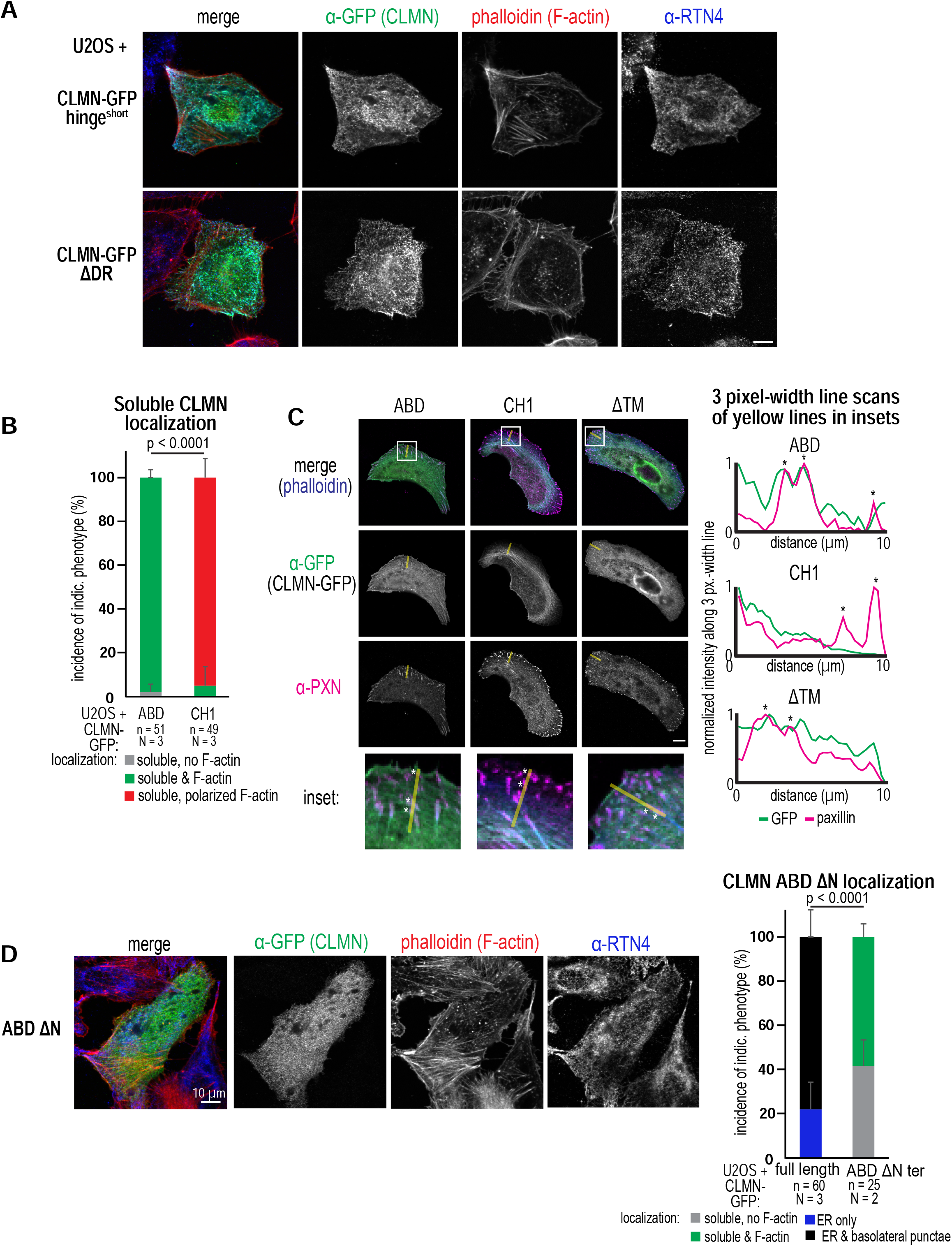
associated with Figure 3: Structure-function analysis of CLMN. A) Laser scanning confocal images of U2OS cells transiently expressing indicated vectors before fixation and labeling with indicated antibodies/dyes. Scale bar = 10 µm. B) Plot of incidences of localization of transiently-expressed CLMN-GFP vectors to the indicated compartments. ABD = actin-binding domain; CH1 = CH1 domain only. n = number of cells within N experimental repeats. Replicate means ± SDs shown. P value = Fisher’s exact test of pooled incidences. C) Laser scanning confocal images of U2OS cells transiently expressing indicated CLMN-GFP truncation constructs and fixed and immunolabeled with indicated antibodies/dyes. Lines indicate line scan areas, and asterisks indicate paxillin foci. Scale bar = 10 µm. Right: 3 pixel-width line scans of normalized fluorescence intensity (normalized to minima/maxima) along yellow lines in images. Asterisks indicate features from images. D) Left: laser scanning confocal images of U2OS cells transiently expressing indicated CLMN-GFP truncation constructs and fixed and immunolabeled with indicated antibodies/dyes. Right: plot of incidences of localization of transiently-expressed CLMN-GFP vectors to the indicated compartments. ABD ΔN ter = actin-binding domain without N terminus. n = number of cells within N experimental repeats. Replicate means ± SDs shown. Full length data is also shown in Figure 3B (left). P value = Fisher’s exact test of pooled incidences.

**Supplementary Figure 5.**
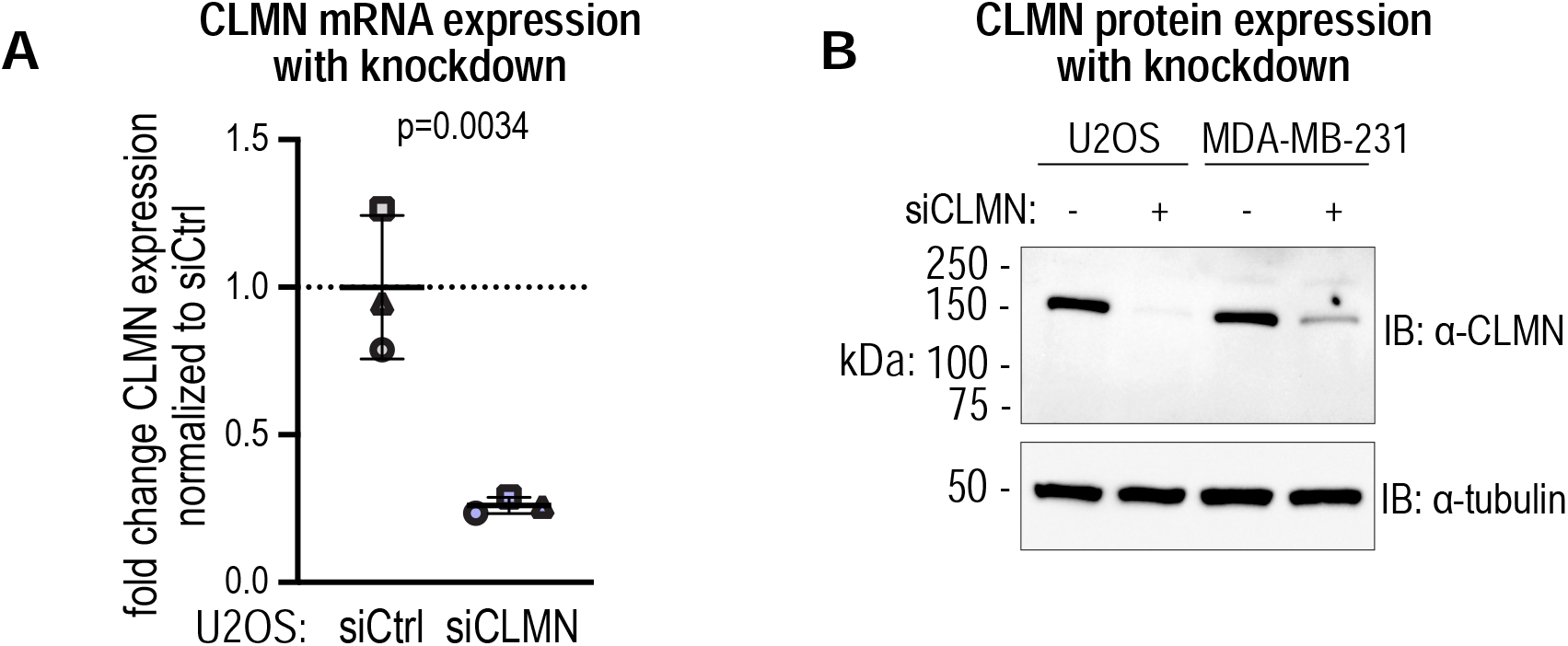
associated with Figure 4: Verification of CLMN knockdowns. A) Plot of fold change of CLMN mRNA expression normalized to siCtrl. 36B4 was used as a loading control. Replicate means ± SDs from N = 3 experimental repeats shown. P value = paired t test of replicate values. B) Immunoblot of whole-cell lysates from indicated cell lines treated with control or CLMN siRNA and immunolabeled with indicated antibodies. Representative image from N = 3 experimental replicates per cell line.

**Supplementary Figure 6.**
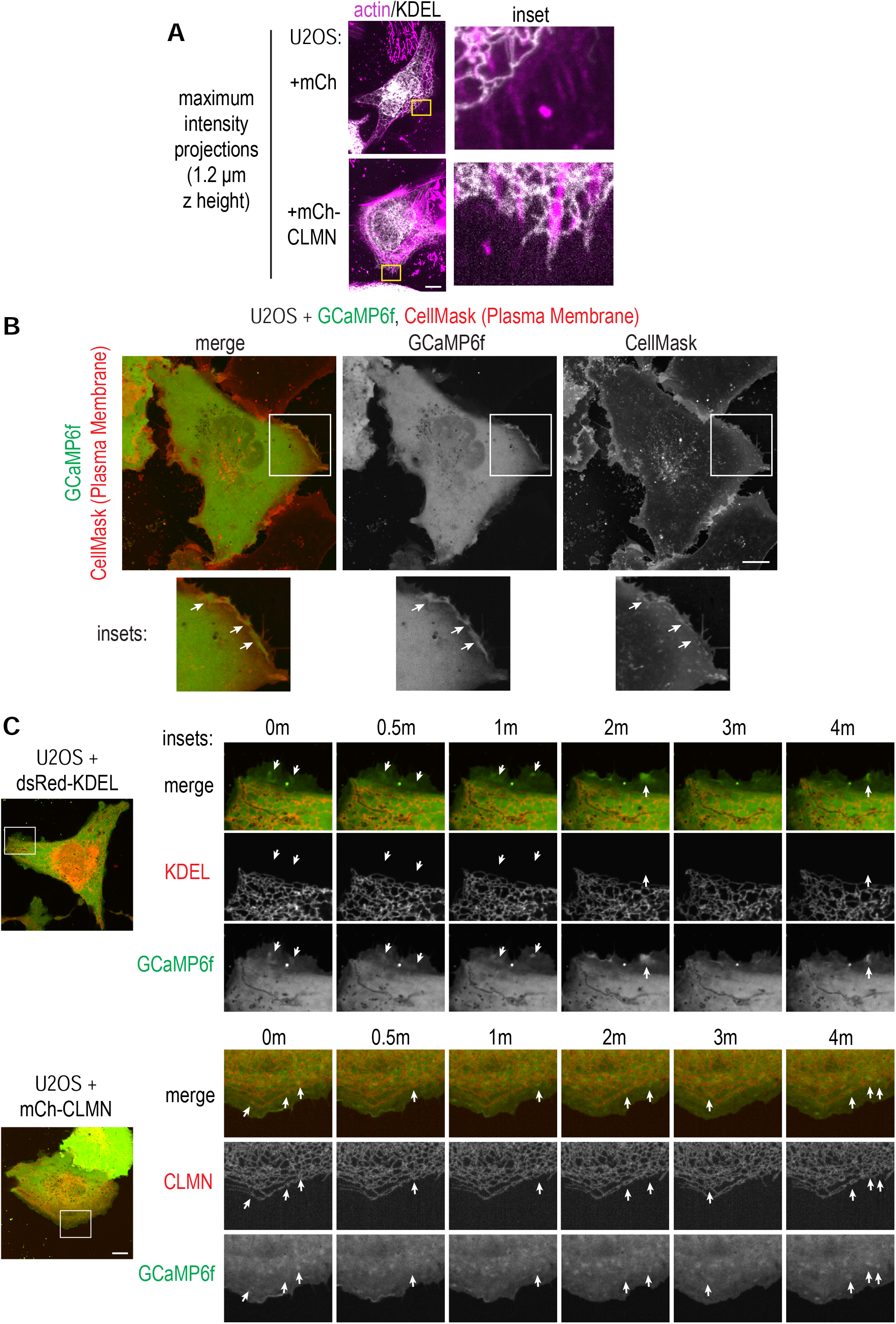
associated with Figure 6: Controls for calcium dynamics in U2OS cells. A) Maximum-intensity projections of live spinning disk confocal images of U2OS cells transiently expressing the indicated vectors and labeled with SiR-actin. Scale bar = 10 µm. Images are maximum-intensity z projections of the same cells depicted in Figure 6A. B) Live spinning disk confocal images of U2OS cells transiently expressing GCaMP6f and labeled with CellMask Orange Plasma Membrane Stain. Arrows indicate GCaMP6f foci that are not associated with a concomitant increase in plasma membrane stain labeling. Scale bar = 10 µm. C) Live spinning disk confocal time lapse images of U2OS cells transiently expressing the indicated vectors. Images are from Figure 6C, with single channel images additionally shown in this figure. Scale bar = 10 µm. Arrows point to GCaMP6f foci.

**Video S1: Localization of CLMN at the cell basolateral surface, pertaining to Figure 2**

Spinning disk confocal time lapse images of CLMN-GFP (green), BFP-KDEL (blue), and SiR-actin (magenta) in a U2OS cell. Image stills are in Figure 2B. Time stamp = (min:sec). Scale bar = 10 µm.

**Video S2: Migration of MDA-MB-231 cells, pertaining to Figure 4**

Phase contrast time lapse images of MDA-MB-231 cells treated as indicated and plated at low density. Image stills are in Figure 4E. Time stamp = (min:sec). Scale bar = 100 µm.

**Video S3: GCaMP6f Ca^2+^ burst foci with CLMN depletion, pertaining to Figure 6**

Spinning disk confocal time lapse images of GCaMP6f in U2OS cells treated as indicated. Image stills are in Figure 6E. Time stamp = (min:sec). Scale bar = 5 µm. White arrows indicate GCaMP burst foci in siControl cells; cyan arrows indicate GCaMP burst foci in CLMN-depleted cells.

**Table S1: Top hits for each Marker Protein used to generate Venn Diagram, pertaining to Figure 1**

Filtered proteomics data from the indicated unmodified and Marker Protein-TurboID-expressing U2OS cell lines briefly treated with biotin and processed (see Methods). Shown are lists of proteins for which fold change > 1 and p < 0.05 for each Marker Protein dataset compared to the un-expressing control (see Methods). Gene names are taken from UniProt.

