## Supplementary material for "Spatial proteomics of ER tubules reveals CLMN, an ER-actin tether at focal adhesions that promotes cell migration": Materials and Methods

### RESOURCE AVAILABILITY

#### Lead Contact

#### Materials Availbility

Unique materials generated in this study are available from the Lead Contact without restriction.

#### Data and code availability

- All original data reported in this study are available from the lead contact upon request.
- All code generated in this study are available from the lead contact upon request.
- All additional information required to re-analyze the data reported in this paper is available from the lead contact upon request.

### EXPERIMENTAL MODEL AND STUDY PARTICIPANT DETAILS

*Tissue Culture of Mammalian Cell Lines*

U2OS, MDA-MB-231, and COS-7 cells and derived cell lines (see Key Resource Table) were obtained from the source specified. Cell lines were grown in 37°C/5% $CO_2$ incubators in media consisting of DMEM high glucose supplemented with 10% heat-inactivated fetal bovine serum, 1% penicillin-streptomycin, and 25 mM HEPES. MDA-MB-231 cell media was additionally supplemented with 1% GlutaMAX Supplement. Cells were tested for mycoplasma upon initial thaw and upon generation of new cell lines using a standard PCR protocol with the mycoplasma-specific oligonucleotides (5'-CCGCGGTAATACATAGGTCGC-3', 5'- CACCATCTGTCACTCTGTTAACC-3'). Cell lines were used before passage 30.

*CRISPR-Cas9 endogenous tagging*

Guide RNA sequences used for gene editing were obtained from the source specified or were designed using the online tool CRISPOR (crispor.tefor.net)[1]. Guide sequences were (PAM sequences in bold): CANX (TCCTAGGACCACTCTTGCAG), RTN4 (GATTATACGGGGGAGGGTCA), CLIMP-63 (GGTGGGCACCCTTCTCCGAG), and LBR (GTGAAGTGGTAAGAGGTCGA). Guide sequences were obtained from the start or stop codon of each gene to edit +/- 100 bp. The guide sequences were synthesized as an oligo pair with BbsI overhangs and were phosphorylated with T4 Polynucleotide Kinase (NEB M0201) in T4 Ligase buffer, annealed by heating to 95C and cooled to 23C. Annealed oligo pairs were cloned into PX459V2.0-HypaCas9[2] digested with BbsI and dephosphorylated using Antarctic Phosphatase.

CRISPR homology-directed repair templates for RTN4, CANX, and LBR were made by amplifying mNeonGreen from pLVX-CMV100-mNeonGreen-Actin-C35[3] and TurboID from V5-TurboID-NES_pCDNA3[4] (Addgene 107169) and combining with synthesized homology repair arms.

Homology repair arms were designed using 800 base pairs upstream and 800 base pairs downstream of the start or stop codons of the genes to be tagged. Sequences were obtained from NIH Gene. Up to 6 base pairs corresponding to PAM sites of potential guide RNA sequences were silently mutated to avoid repair template cutting by Cas9. Homology repair arms were synthesized by Twist Biosciences or by Integrated DNA Technologies. Homology repair arms were synthesized with linkers such that the target gene and TurboID were separated by an AGSAGSAAGSGEF linker, and TurboID and mNeonGreen were separated by a GGSGGGGSGGGGS linker. The homology repair arms, TurboID, and mNeonGreen were assembled into an EGFP-N2 backbone lacking the CMV and GFP sequences using Gibson Assembly.

For CLIMP-63, the mNeonGreen-TurboID insert from the LBR repair template was amplified with overhangs corresponding to 37 bp upstream and downstream of the CLIMP-63 start codon as in[5], purified using the Promega Wizard PCR purification kit, and used as a PCR-based repair template.

For gene editing, guide RNA and repair template plasmids were dually transfected into cells using Lipofectamine 3000 according to the manufacturer's protocol. After 24 hours of transfection, cells were treated with 2-3 μg/ml of puromycin for 24 hours. Cells were then grown up and sorted once by mNeonGreen fluorescence to obtain enriched populations of knock-in cells using a BD FACSAria I. mNeonGreen-TurboID-CLIMP-63$^{EN}$ cells were sorted twice to enrich for mNeonGreen-positive cells, and RTN4$^{EN}$-TurboID-mNeonGreen cells were processed to obtain a clonal knock-in cell line due to an off-target population. Full-length integration was determined by Western blotting of mNeonGreen in whole-cell lysates from each cell line. For LBR$^{EN}$, full-length integration was additionally confirmed by PCR of a genomic region encompassing the mNeonGreen-TurboID-LBR repair template using genomic DNA extracted from U2OS mNeonGreen-TurboID-LBR$^{EN}$ using a PureLink gDNA extraction kit.

*Stable cell line generation*

To generate a U2OS cell line stably expressing mNeonGreen-TurboID-Nesprin4, the coding sequence of Nesprin 4 was synthesized with Twist Biosciences and cloned into EGFP-N2 with the GFP insert swapped for mNeonGreen-TurboID-Nesprin 4. U2OS cells were transfected with the mNeonGreen-TurboID-Nesprin 4 vector using Lipofectamine 3000; after 24 hours of transfection, cells were treated with 400 ug/ml G418 for 2 weeks, then cells were processed to obtain a clonal cell line. Nesprin 4 full-length integration was confirmed by PCR of the mNeonGreen-TurboID-Nesprin 4 insert using genomic DNA extracted from the stable cell line.

*Clonal cell line generation*

Cells were plated at <10 cells/ml in a 10 cm dish and grown for 2-3 weeks until colonies of 50-100 cells were easily visible. Colonies were marked with a marker underneath the dish at 5-10x magnification. Cells were washed 1x with PBS, then sterile cloning discs were dipped in 0.25% trypsin-EDTA and laid onto the colonies using sterile tweezers. After 2-5 min of trypsinization, cloning discs were lifted from the dish and plated into 12-well plates containing 2 mls of media. Cells were grown to confluence and then imaged/genotyped.

### METHOD DETAILS

*DNA and RNAi Transfection*

DNA transfections were performed using Lipofectamine 3000 in Opti-MEM Reduced Serum Medium. For transfection mixes, each 1 ml of final mixture in growth medium contained 100 µl of Opti-MEM, 0.5-1 µg of DNA, 2 µl of P3000, and 1.5 µl of Lipofectamine 3000. DNA/P3000 and Lipofectamine 3000 were diluted in Opti-MEM and incubated for 5 minutes. The DNA/P3000 mixture was added dropwise to the Lipofectamine mixture and incubated for 20 minutes before adding dropwise to wells of cells containing growth media. Media was exchanged after 6-24 hours. RNAi was performed using Lipofectamine RNAiMAX. For CLMN knockdowns, CLMN Silencer Select siRNA or Silencer Select negative control siRNA #1 were diluted in 250 µl Opti-MEM and then into full media (with a final concentration of 30 nM), with Lipofectamine RNAiAX used at a concentration of 3 µl/ml. RNA diluted in Opti-MEM was added dropwise to Lipofectamine RNAiMAX diluted in Opti-MEM and incubated for 5 minutes before adding to cells in growth media. Media was changed after 24 hours, and total RNAi time was 48 hours. For experiments using dual RNAi/DNA transfection, RNAi was performed for 48 hours, and DNA transfection was performed the following day for 24 hours.

*Expression plasmid generation*

CLMN was amplified from a cDNA clone from The Ultimate ORF Lite human cDNA collection and cloned into EGFP-N2 or mCherry-C1 using Gibson Assembly. CLMN truncation mutants were generated by PCR amplification, gene fragment synthesis (Integrated DNA Technologies), or annealing short oligonucleotides corresponding to regions before and after deletions, which were subsequently cloned into EGFP-N2 using Gibson Assembly. CLMN plasmids were verified by nanopore whole-plasmid sequencing by Plasmidsaurus or Eurofins Genomics. CLMN vector mutations corresponded to the following amino acid positions: ΔCH1 deletion of a.a. 32-139, ΔCH2 deletion of a.a. 187-291, ΔCH1CH2 deletion of a.a. 32-291, CLMN-hinge$^{short}$ deletion of a.a. 148-178, ΔTM deletion of a.a. 977-994, ΔC ter deletion of a.a. 977-1002, ΔDR deletion of a.a. 389-911, CLMN-TM$^{extend}$ added ILLF after a.a. 994, CLMN-TM$^{Sec61\beta}$ swapped 977-994 of CLMN for 21 a.a. transmembrane segment of Sec61β, ABD retained only a.a. 1-300, ADB ΔN ter retained only a.a. 32-291, CH1 retained only a.a. 1-145.

*Quantitative Real-Time PCR*

RNA was extracted from cultured cells using Trizol-chloroform extraction. Cell pellets were washed with PBS and centrifuged at 400xg for 5 minutes. 0.5 ml Trizol was added and pipetted up and down to mix, then incubated for 5 minutes at room temperature. Chloroform was added at 200 µl per ml, and extraction mixtures were shaken for 15 seconds before incubating at room temperature for 3 minutes and centrifugation at 12000xg for 15 minutes at 4°C. The colorless upper aqueous fraction was pipetted into a new 1.5 ml tube, and 500 µl of isopropanol per 500 µl was added and the tube inverted 4x before being incubated at room temperature for 10 minutes and centrifuged at 12000xg for 10 minutes at 4°C. The supernatant was removed, and the resulting pellet was washed with 75% ethanol, vortexed, centrifuged at 7500xg for 10 minutes at 4°C, and the supernatant was aspirated. The pellet was dried, resuspended in nuclease-free water, and incubated at 55°C for 10 minutes before being stored on ice. To synthesize cDNA, 1 µg of RNA was combined into a 20 µl 1x mixture of iScript Reverse Transcription Supermix for qRT-PCR. Cycler conditions were 5 minutes at 25°C, 20 minutes at 46°C, 5 minutes at 95°C, then cold hold. For qRT-PCR, cDNA was diluted to 500 ng/µl, and reaction mixes were set up using SsoAdvanced Universal SYBR Green Supermix and 1.6 µM primers with 1 µg of cDNA per 20 µl reaction. qRT-PCR was performed on a BioRad CFX96 Real-Time system, and mRNA expression data was normalized to expression of 36B4 (see Key Resources Table).

*Immunoblot*

Protein lysates were extracted in RIPA buffer (50 mM Tris pH 7.4, 150 mM NaCl, 0.1% SDS, 0.5% sodium deoxycholate, 1% Triton X-100) containing Halt protease or protease/phosphatase inhibitor. Lysates were incubated on ice for 20 minutes and pushed through a 23G needle 24x, then centrifuged at 21000xg for 10 minutes at 4°C. Lysate concentrations were quantified using the Pierce Detergent-Compatible Bradford assay, and 10-20 µg of protein was loaded into Mini-PROTEAN Precast TGX gels (Bio-Rad 45610) and ran at 80-100V. Protein was wet-transferred to 0.45 µm supported nitrocellulose or PVDF membranes. Membranes were blocked in 5% blotting grade blocker in TBS for 1 hour, then incubated in primary/secondary antibody solutions in 5%

blotting grade blocker in TBS-Tween for 1 hour followed by 3 5-10 minute washes in TBS-Tween after each antibody incubation step. Membranes were incubated in Clarity ECL or Clarity Max ECL reagent (Bio-Rad 1705061, 1705062) for 1-2 minutes and imaged in a Bio-Rad GelDoc imager. Exposures were set to the maximum exposure time before saturated pixels began to appear.

Antibody dilutions: α-mNeonGreen 1:500, Streptavidin-HRP 1:3000-1:5000, α-tubulin α 1:5000, α-CLMN 1:100, α-GAPDH 1:2000

*Proteomics*

2 days prior to harvesting cell lysates for proteomic analysis, cells were plated at 5 x $10^6$ cells into 4 x T125 flasks each. The next day, media was exchanged for DMEM with 25 mM HEPES and 100 µM BSA. The following day, cells were treated with 50 µM biotin in DMEM for 10 minutes at 37°C before putting flasks on ice. Cells were washed 3x with cold PBS, and cells were collected by scraping. Cells were centrifuged at 500xg for 5 minutes at 4°C, and pellets were flash frozen in liquid nitrogen and stored at -80°C. When all samples were processed, pellets were thawed on ice and resuspended in RIPA buffer (see Immunoblot section) containing Halt protease inhibitor. Lysates were incubated on ice for 40 minutes and pushed through a 25G needle 12x. Lysates were centrifuged at 13,000xg for 10 minutes at 4°C; the supernatant was removed to new pre-chilled tubes. 3500 MWCO Slide-A-Lyzer dialysis cassettes were soaked in RIPA buffer. To remove excess biotin, samples were dialyzed in cassettes overnight at 4°C with gentle stirring, with some sample saved as undialyzed input. After dialysis, Halt protease inhibitor was added to a final concentration of 1x; Protein was quantified by Bradford assay. Enrichment of biotinylated proteins proceeded as in[6]. Streptavidin-conjugated magnetic beads were washed twice with RIPA buffer. 0.25-0.75 mg of protein was added to 100 µl beads, with some sample saved as dialyzed input. Samples were incubated on beads overnight with rotation at 4°C, then centrifuged briefly at 500xg to collect the liquid. Magnetic beads were washed with MS-compatible RIPA buffer (50 mM Tris pH 7.4, 150 mM NaCl, 0.5% sodium deoxycholate) 2x, then incubated in 1 M KCl for 2 minutes, then incubated in 0.1 M $Na_2CO_3$ for 10 seconds, then incubated in 2 M urea in 10 mM Tris-HCl pH 8 for 10 seconds, then washed with MS-compatible RIPA buffer, then resuspended in MS-

compatible RIPA buffer with 1x Halt protease inhibitor. Sample protein was eluted by boiling in 3x loading buffer (made from 6x loading buffer: 0.33 M Tris pH 6.8, 34% glycerol, 10% SDS, 0.09% DTT, 0.12% bromophenol blue) with 2mM biotin and 20 mM DTT at 95°C for 10 minutes. Samples were centrifuged at 500xg for 30 seconds, then pelleted, and eluates were collected and combined with 6x loading buffer and boiled at 95°C for 10 minutes, with some eluates saved for quality control analysis. Eluates were flash frozen in $LN_2$ until ready to process. Quality control analysis was done by immunoblotting for Streptavidin-HRP (1:3000 antibody dilution, 30 minutes incubation in 3% BSA in TBS after membrane blocking) in undialyzed input, dialyzed input, and eluate samples, and by using the Pierce Silver Stain kit to visualize whole protein in undialyzed input, dialyzed input, and eluate samples.

For trypsin digestion and label-free MS, eluate samples were thawed at room temperature, boiled at 95°C for 2 minutes, and centrifuged. Samples were loaded into 10% TGX Mini-PROTEAN 50 µl well gels and run for 0.5 cm into the gels. Gels were stained with 0.1% Coomassie BB R250 in 10% acetic acid and 50% methanol for 15 minutes, then destained for 1.5 hours in 10% acetic acid and 50% methanol with >2 solvent changes; destain was then exchanged for milliQ $H_2O$. Protein bands were cut from each lane, diced, and stored at 4°C for delivery to the Proteomics Core Facility at UT Southwestern Medical Center.

Samples were digested overnight with trypsin (Pierce) following reduction and alkylation with DTT and iodoacetamide (Sigma–Aldrich). The samples then underwent solid-phase extraction cleanup with an Oasis HLB plate (Waters) and the resulting samples were injected onto an Orbitrap Fusion Lumos mass spectrometer coupled to an Ultimate 3000 RSLC-Nano liquid chromatography system. Samples were injected onto a 75 um i.d., 75-cm long EasySpray column (Thermo) and eluted with a gradient from 0-28% buffer B over 90 min. Buffer A contained 2% (v/v) ACN and 0.1% formic acid in water, and buffer B contained 80% (v/v) ACN, 10% (v/v) trifluoroethanol, and 0.1% formic acid in water. The mass spectrometer operated in positive ion mode with a source voltage of 2.0 kV and an ion transfer tube temperature of 275 °C. MS scans were acquired at 120,000 resolution in the Orbitrap and up to 10 MS/MS spectra were obtained in the ion trap for each full spectrum acquired using higher-energy collisional dissociation (HCD) for ions with charges 2-7. Dynamic exclusion was set for 25 s after an ion was selected for fragmentation.

Raw MS data files were analyzed using Proteome Discoverer v.2.4 SP1 (Thermo), with peptide identification performed using Sequest HT searching against the reviewed human protein databases from UniProt as well as the sequences of TurboID and mNeonGreen. Fragment and precursor tolerances of 10 ppm and 0.6 Da were specified, and three missed cleavages were allowed. Carbamidomethylation of Cys was set as a fixed modification, with oxidation of Met set as a variable modification. The false-discovery rate (FDR) cutoff was 1% for all peptides. Proteins that were identified with >1 peptide were considered for analysis. If more than one protein in a group had the same score, an equal number of peptide spectrum matches, and an equal number of peptides, the protein with the longest sequence was designated as the master protein.

For each condition, protein abundances were normalized to the average total peptide abundance for the unmodified biotin-treated U2OS samples. Sample values were averaged across experimental repeats and expressed as a fold change over the unmodified biotin-treated U2OS sample average. 2-tailed t tests were used to obtain p values between each condition and the unmodified U2OS control; protein lists were sorted in R to grab proteins with a fold change > 1 and $p < 0.05$; these final lists were used to make a 5-point Venn diagram using the VennDiagram package in R. Volcano plots of $\log_2$ fold change and $-\log_{10}$ p values were used to identify key top hits for each protein, including CLMN in the RTN4 dataset.

*Immunofluorescence*

Cells were plated onto No. 1.5 acid-washed coverslips or in Nunc Labtek glass-bottom imaging dishes (Thermo Scientific 155409). For experiments imaging cells migrating toward a wound area (i.e. experiments quantifying paxillin foci), a hatch-shaped scratch was inflicted on the culture surface using a p10 pipette tip, then cells were washed 2x with PBS and incubated in growth media for 2 hours prior to fixation. For imaging biotinylated proteins, cells were treated with serum-free media with 100 μM BSA for 16-

24 hrs, then treated with 50 μM biotin in serum-free media for 10 minutes prior to washing 2x with PBS on ice. For all other immunofluorescence applications, cells were washed with room-temperature or 37°C PBS 1x prior to fixation. All fixations were performed with 4% paraformaldehyde in PBS for 15 minutes at room temperature. Cells were permeabilized with 0.5% Triton X-100 in PBS for 5 minutes, then washed 3x with PBS. Cells were blocked with 3% BSA in PBS for 30 minutes, then incubated in primary/secondary antibody solutions in 3% BSA in PBS for 1 hour followed by 3 5-minute washes in PBS after each antibody incubation step. SiR-actin, phalloidin, or streptavidin conjugates were included in primary incubation steps when indicated. Cells were mounted onto glass slides or mounted in-well with 90% glycerol in 20 mM Tris pH 8.7 supplemented with 0.5% N-propyl gallate. Coverslips were sealed with clear nail polish or CoverGrip sealant (Biotium 23005).

Antibody/dye dilutions: Alexa Fluor 546 Phalloidin 1:400, SiR-actin 1:1000, Alexa Fluor 647-streptavidin 1:500, α-mNeonGreen 1:500-1:1000, α-CANX 1:500, α-RTN4 1:200-1:500, α-GFP 1:500-1:1000, α-paxillin 1:250, Alexa Fluor 488 Donkey anti-mouse and goat anti-rabbit 1:500, Rhodamine RedX Donkey anti-rabbit and goat anti-mouse 1:500, Alexa Fluor 647 Donkey anti-rabbit and goat anti-mouse 1:500-1:2000

*Microscopy*

Imaging fixed samples was performed on a Zeiss LSM 780 or LSM 880 inverted laser scanning confocal microscopes with 405, 458, 488, 514, 561, 594, and 633 nm laser lines and 10x-63x objectives, Live imaging was performed on a Nikon TiE inverted microscope with a Yokogawa CSU-W1 spinning disk with SoRa module, 405, 445, 488, 514, 561, and 640 nm laser lines, 4x-100x objectives, a Hamamatsu ORCA-Fusion sCMOS camera, and Oko Labs environmental control with a stagetop incubator for 37°C/5% $CO_2$ imaging.

*Live-cell imaging*

For live cell imaging, cells were plated in either 8-well Nunc Labtek glass-bottom dishes, μ-slide 8-well glass bottom dishes (ibidi 80827), or 35 mm glass-bottom dishes. For non-$CO_2$-controlled short-term live cell imaging, cells were imaged in live imaging media consisting of 140mM NaCl, 2.5 mM KCl, 1.8 mM $CaCl_2$, 1.0 mM $MgCl_2$, 20mM HEPES, pH 7.4 supplemented with 15 mM glucose. For time-lapse imaging that is $CO_2$-controlled, cells were imaged in Fluorobrite DMEM (Gibco A1896701). MDA-MB-231 cells were imaged in Fluorobrite DMEM supplemented with 10% FBS and 1x GlutaMAX.

For SiR-actin labeling, cells were incubated in 1:200-1:1000 SiR-actin in live imaging media for 15-45 minutes prior to imaging. For CellMask Orange labeling, cells were incubated in 1:1000 CellMask in live imaging media for 7 minutes, then washed 3x with media, then imaged in fresh live imaging media within 60 minutes.

*Cell migration assays*

For scratch assays in U2OS, cells were treated with 30 nM control or CLMN siRNA for 24 hours before plating. Cells were plated at 300,000 cells per well in a 12-well dish to be 100% confluent the following day. After 48 h of RNAi and 24 h after plating, cross shapes were scratched onto the culture surfaces using a p10 pipette tip, and the shapes were marked underneath the dishes using a marker at 10x magnification. Cells were washed 1x with PBS, and media was replaced with growth media containing 1:1000 mitomycin to inhibit proliferation. Cultures were imaged at 0 h and 10 h after wounding using a Bio-Rad ZOE Fluorescent Cell Imager set to the epifluorescence phase contrast setting. Cultures were imaged at the pre-marked locations to ensure the same location was imaged at each time point. To quantify wound size, the edges of the wound in each image were blindly manually traced using the polygon tool in ImageJ and area measured. Wound sizes at each point were the average of measurements from 2 fields of view. Wound sizes were normalized to the average of the control at time 0 h.

For single cell migration assays in MDA-MB-231 (Gau & Roy, 2020), cells were treated with 30 nM control or CLMN siRNA for 24 hours before plating or were transfected with the indicated DNA vectors 24 hours before plating. Cells were plated at 10,000 cells per well in an ibidi 8-well imaging dish. The following day, cells were imaged with 5-7 fields of view taken from each well every 1 minute for 120 minutes using phase contrast at 10x magnification using multi-point revisiting. Transfected cells were imaged in the 488/GFP channel just before starting the time lapse to correlate which cells were expressing the indicated protein. To measure average cell velocity, single cells were chosen for analysis and analyzed as in[7]. The ImageJ Manual Tracker tool was used to track the position of single cells over time, using the nucleus as the center of the cell. Positions over time were converted to average velocity using the formula

$$v = \frac{\sum \sqrt{(x_2 - x_1)^2 + (y_2 - y_1)^2}}{n}$$

Where v is average velocity, $x_2$ and $x_1$ correspond to x positions at time points 2 and 1, $y_2$ and $y_1$ correspond to y positions at time points 2 and 1, and n refers to the total number of time points.

*Image analysis*

Image analysis was performed using FIJI/ImageJ[8]. All images with multiple markers/colors in the same image are composite images (multiple markers shown as different colors within composite images). All data quantifying incidence of localization, scoring of events, or phenotypic scoring were performed blindly; images were blinded using the Filename_Randomizer macro in ImageJ modified to accommodate .nd2 files. Representative intensity plot profiles/line scans were generated using the Plot Profile function in ImageJ using straight lines with pixel widths and lengths indicated in each figure. For quantification using "cell peripheral edge area" as a parameter, the cell peripheral edge area was delineated using the fluorescence drop-off of soluble GCaMP6f at the cell peripheral edge; edge areas were pre-defined before analysis using manual ROI drawing and area quantification. For quantification of GCaMP6f foci occurring at the cell peripheral edge relative to ER localization, foci occurring at the plasma membrane were excluded from analysis.

Paxillin foci were quantified using Weka machine learning segmentation[9] in ImageJ. Classifiers were made by training the model with 400 events for background and 400 events for paxillin foci. The same classifier was used within all conditions of each experimental repeat, but classifiers were re-made for some experimental repeats to account for differences in immunofluorescence staining across repeats. Classifiers were applied to single z sections corresponding to the coverslip surface of each image, then classified images were converted to 8-bit, then images were converted to a mask and subject to watershed to split up converging foci. ROIs corresponding to the entire cell border for cells at the migrating front (when applicable) were manually traced using the phalloidin channel of each image. For a cell area ROI, Particle Analysis (ImageJ Analyze Particles function) was performed to quantify the number and size of paxillin foci per cell area.

### QUANTIFICATION AND STATISTICAL ANALYSIS

GraphPad Prism 8 and Microsoft Excel were used for all statistical analysis. In imaging experiments wherein phenotypes or localization incidences of individual cells were scored, n refers to individual cells unless otherwise stated; for all, N refer to number of experimental repeats. Superplot format[10] and paired t test/ANOVA of replicate means were utilized in experiments plotting continuous data in which n > 10 per experimental repeat was feasible. Discrete data with 2 categories were plotted with separate shapes or colors indicating separate experimental repeats. Statistical tests, sample sizes, and p values are reported in figures/figure legends. Sample size calculations for discrete data with 2 categories were performed using the online tool (https://clincalc.com/stats/samplesize.aspx) to determine the adequate sample size for number of cells to analyze for sufficient (80%) power. Continuous data were tested for normality (and whether to use parametric or non-parametric tests) using the Shapiro-Wilk test.

### KEY RESOURCES TABLE

| REAGENT or RESOURCE | SOURCE | IDENTIFIER |
|---|---|---|
| Antibodies | | |
| Rabbit anti-calnexin | Abcam ab22595 | RRID:AB_2069006 |
| Mouse anti-mNeonGreen | Proteintech 502256466 | RRID:AB_2827566 |
| Mouse anti-RTN4 | Santa Cruz sc-271878 | RRID:AB_10709573 |
| Rabbit anti-GFP | Abcam ab290 | RRID:AB_303395 |
| Rabbit anti-CLMN | Atlas Antibodies HPA012634 | RRID:AB_1846973 |

| | | |
|---|---|---|
| Mouse anti-paxillin | BD Biosciences 610051 | RRID:AB_397463 |
| Rat anti-tubulin | Abcam ab6160 | RRID:AB_305328 |
| Rabbit anti-GAPDH | Abcam ab9485 | RRID:AB_307275 |
| Goat anti-Rabbit HRP | Abcam ab6721 | RRID:AB_955447 |
| Rabbit anti-mouse HRP | Abcam ab6728 | RRID:AB_955440 |
| Goat anti-Rat HRP | Abcam ab205720 | RRID:AB_2941939 |
| Alexa Fluor 488 goat anti-rabbit | Thermo Fisher A11034 | RRID:AB_2576217 |
| Alexa Fluor 488 Donkey anti-mouse | Thermo Fisher A21202 | RRID:AB_141607 |
| Rhodamine RedX Donkey anti-rabbit | Jackson Immuno 711-295-152 | RRID:AB_2340613 |
| Rhodamine RedX goat anti-mouse | Jackson Immuno 115-295-146 | RRID:AB_2338766 |
| Alexa Fluor 647 Donkey anti-rabbit | Thermo Fisher A32795 | RRID:AB_2762835 |
| Alexa Fluor 647 Goat anti-mouse | Thermo Fisher A3272872 | RRID:AB_2633277 |
| Chemicals, peptides, and recombinant proteins | | |
| Puromycin | Gibco A11138 | |
| G418 | Invivogen ant-gn-1 | |
| 0.25% trypsin-EDTA | Sigma T4049 | |
| DMEM high glucose | Gibco 11965 | |
| Heat-inactivated FBS | Sigma F4135 | |
| Penicillin-streptomycin | Sigma P4333 | |
| 25 mM HEPES | Sigma H0887 | |
| GlutaMAX supplement | Thermo Fisher 35050061 | |
| T4 Polynucleotide Kinase | NEB M0201 | |
| T4 Ligase buffer | NEB M0202 | |
| BbsI | NEB R0539 | |
| Antarctic Phosphatase | NEB M0289 | |
| Gibson Assembly Mastermix | NEB E2611 | |
| Lipofectamine 3000 | Thermo Fisher L3000 | |
| Lipofectamine RNAiMAX | Invitrogen 13778 | |
| Opti-MEM | Gibco 11058 | |
| Trizol | Life Technologies 15596018 | |
| iScript Reverse Transcription Supermix | Bio-Rad 1708841 | |

| | | |
|---|---|---|
| SsoAdvanced Universal SYBR Green Supermix | Bio-Rad 1725274 | |
| Halt protease inhibitor | Thermo Scientific 78429 | |
| Halt protease/phosphatase inhibitor | Thermo Scientific 78440 | |
| Mini-PROTEAN Precast TGX gel | Bio-Rad 45610 | |
| blotting grade blocker | Bio-Rad 1706404 | |
| Clarity ECL reagent | Bio-Rad 1705061 | |
| Clarity Max ECL reagent | Bio-Rad 1705062 | |
| Coomassie Brilliant Blue R250 | Fisher BP101-50 | |
| Fluorobrite DMEM | Gibco A1896701 | |
| mitomycin | Millipore Sigma M5353 | |
| Critical commercial assays | | |
| Pierce Detergent-Compatible Bradford assay | Thermo Scientific 23246 | |
| Pierce Silver Stain kit | Thermo Scientific 24612 | |
| Experimental models: Cell lines | | |
| U2OS | ATCC | RRID:CVCL_0042 |
| COS-7 | Jonathan Friedman lab, UT Southwestern Medical Center | RRID:CVCL_0224 |
| MDA-MB-231 | ATCC | RRID:CVCL_0062 |
| U2OS CANX$^{EN}$-TurboID-mNeonGreen | This study | |
| U2OS RTN4$^{EN}$-TurboID-mNeonGreen | This study | |
| U2OS mNeonGreen-TurboID-CLIMP63$^{EN}$ | This study | |
| U2OS mNeonGreen-TurboID-LBR$^{EN}$ | This study | |
| U2OS stably expressing mNeonGreen-TurboID-Nesprin4 | This study | |
| Oligonucleotides | | |
| Mycoplasma testing FWD - CCGCGGTAATACATAGGTCGC | Sigma-Aldrich #MP0025 | |
| Mycoplasma testing REV - CACCATCTGTCACTCTGTTAACC | Sigma-Aldrich #MP0025 | |
| CLMN Silencer Select siRNA | Thermo Fisher 4392420, siRNA ID s36333 | |
| Silencer Select negative control siRNA #1 | Thermo Fisher 4390843 | |

| | | |
|---|---|---|
| CLMN qPCR primer FWD<br>AGCGACTACAGCATTCCTTCC | This study | |
| CLMN qPCR primer REV<br>GCTGTTGGCCTTCCTGGTTA | This study | |
| 36B4 qPCR primer FWD<br>ACATCCGAGTGCAGAACCTG | Neuhaus et al., 2011[11] | |
| 36B4 qPCR primer REV<br>AGAGGTGGGTTGCACTTTTCTA | Neuhaus et al., 2011[11] | |
| $LBR^{EN}$ genotyping primer FWD<br>GGAAATGTGATTCCGCTGGTC | This study | |
| $LBR^{EN}$ genotyping primer REV<br>ATGGGTCCTAGAAATGCTCGT | This study | |
| Nesprin 4 genotyping primer FWD (mNeonGreen FWD)<br>ATGGTGAGCAAGGGCGAGG | This study | |
| Nesprin 4 genotyping primer REV<br>TCAGACTGGGGGAAGACCA | This study | |
| Recombinant DNA | | |
| pLVX-CMV100-mNeonGreen-Actin-C35 | Mohan et al., 2019[3] | |
| V5-TurboID-NES_pCDNA3 | Addgene 107169 | RRID:Addgene_107169 |
| PX459V2.0-HypaCas9 | Addgene 108294 | RRID:Addgene_108294 |
| BFP-KDEL | Addgene 49150 | RRID:Addgene_49150 |
| Rtn4a-GFP | Addgene 61807 | RRID:Addgene_61807 |
| mCherry-C1 | Clontech | |
| EGFP-N2 | Clontech | |
| dsRed-KDEL | TaKaRa | |
| GCaMP6f | Addgene 40755 | RRID:Addgene_40755 |
| mCh-CLMN | This study | |
| CLMN-GFP | This study | |
| CLMN-GFP si-resistant | This study | |
| CLMN-GFP $TM^{Sec61\beta}$ | This study | |
| CLMN-GFP $TM^{extended}$ | This study | |
| CLMN-GFP si-resistant ΔCH1 | This study | |

| | | |
|---|---|---|
| CLMN-GFP si-resistant ΔCH2 | This study | |
| CLMN-GFP si-resistant ΔCH1CH2 | This study | |
| CLMN-GFP si-resistant hinge$^{short}$ | This study | |
| CLMN-GFP si-resistant ΔTM | This study | |
| CLMN-GFP si-resistant ΔC ter | This study | |
| CLMN-GFP si-resistant ΔCH1CH2 | This study | |
| CLMN-GFP si-resistant ΔIDR | This study | |
| CLMN ABD-GFP | This study | |
| CLMN ABD-GFP ΔN ter | This study | |
| CLMN CH1-GFP | This study | |
| Software and algorithms | | |
| ImageJ | Schindelin et al., 2012[8] | RRID:SCR_003070 |
| GraphPad Prism 10 | GraphPad Software | RRID:SCR_002798 |
| Microsoft Excel | Microsoft, Inc. | RRID:SCR_016137 |
| R | The R Project | RRID:SCR_001905 |
| R Studio | Posit | RRID:SCR_000432 |
| R package Tidyverse | Posit | RRID:SCR_019186 |
| R package VennDiagram | Hanbo Chen (UCLA) | RRID:SCR_002414 |
| ImageJ Filename_Randomizer Macro | Tiago Ferreira (EMBL) | |
| Other | | |
| Sterile cloning discs | Sigma Z374431 | |
| Wizard SV gel and PCR purification kit | Promega A9282 | |
| PureLink Genomic DNA Mini Kit | Invitrogen K182002 | |
| The Ultimate ORF Lite human cDNA collection | Life Technologies | |
| 3500 MWCO Slide-a-Lyzer dialysis cassettes | Thermo Scientific 66330 | |
| Streptavidin-conjugated magnetic beads | Thermo Scientific 88817 | |
| Streptavidin-HRP | Invitrogen 19534 | |
| Nunc Labtek glass-bottom imaging dishes | Thermo Scientific 155409 | |
| CoverGrip sealant | Biotium 23005 | |
| µ-slide 8-well glass bottom dishes | ibidi 80827 | |
| 35 mm glass-bottom dishes | Fisher Scientific D35141.5N | |

| | |
|---|---|
| SiR-actin | Cytoskeleton, Inc. CY-SC001 |
| Alexa Fluor 546 phalloidin | Invitrogen A22283 |
| Alexa Fluor 647 Streptavidin | Invitrogen S32357 |
| CellMask Orange plasma membrane stain | Invitrogen C10045 |
